# Clade-wise alignment integration improves co-evolutionary signals for protein-protein interaction prediction

**DOI:** 10.1101/2023.07.28.550005

**Authors:** Tao Fang, Damian Szklarczyk, Radja Hachilif, Christian von Mering

**Affiliations:** Department of Molecular Life Sciences, University of Zurich, 8057 Zurich, Switzerland; SIB Swiss Institute of Bioinformatics, 1015 Lausanne, Switzerland

## Abstract

**Background:** Protein-protein interactions play essential roles in almost all biological processes. The binding interfaces between interacting proteins impose evolutionary constraints, leading to co-evolutionary signals that have successfully been employed to predict protein interactions from multiple sequence alignments (MSAs). During the construction of MSAs for this purpose, critical choices have to be made: how to ensure the reliable identification of orthologs, how to deal with paralogs, and how to optimally balance the need for large alignments versus sufficient alignment quality.

**Results:** Here, we propose a divide-and-conquer strategy for MSA generation: instead of building a single, large alignment for each protein, multiple distinct alignments are constructed, each covering only a single clade in the tree of life. Co-evolutionary signals are searched separately within these clades, and are only subsequently integrated into a final interaction prediction using machine learning. We find that this strategy markedly improves overall prediction performance, concomitant with better alignment quality. Using the popular DCA algorithm to systematically search pairs of such alignments, a genome-wide all-against-all interaction scan in a bacterial genome is demonstrated.

**Conclusions:** Given the recent successes of AlphaFold in predicting protein-protein interactions at atomic detail, a discover-and-refine approach is proposed: our method could provide a fast and accurate strategy for pre-screening the entire genome, submitting to AlphaFold only promising interaction candidates - thus reducing false positives as well as computation time.

## Introduction

At evolutionary timescales, proteins tend to accumulate changes in their amino acid sequences, whereas their three-dimensional structure remains largely conserved^1, 2^. This is because only a minority of residues are absolutely essential for the correct folding and function. In contrast, other residues are relatively free to change during genetic drift or in response to secondary constraints arising from changed organismic lifestyles (temperature, salinity, etc.)^3–5^. An additional degree of freedom in protein sequence evolution is afforded by “compensatory mutations’’: mutations that would be detrimental on their own can be compensated for by other mutations elsewhere in the protein. Compensatory mutations are often (but not always) proximal to each other in the three-dimensional protein structure. Hence, they create a statistical co-evolution signal that has become an essential ingredient when predicting protein structure from sequences^6–8^.

Coevolution is traditionally used to study proteins’ three-dimensional (3D) structures and functions within the same protein (family). Most prominent computational methods nowadays to detect pairwise co-evolving residues include local statistical methods such as mutual information (MI) based methods^6, 7, 9, 10^ and global statistical methods such as sparse inverse covariance estimation (PSICOV)^11^ and direct coupling analysis (DCA)^12^. While local statistical methods like MI are very efficient, they only consider one residue pair at a time. Therefore it can not distinguish between direct contact and indirect contact. This so-called transitivity problem arises from chains of direct coupling, i.e., two non-contacting residues can show a high coupling score when they both are coupled with third residues^13–17^. Global statistical models such as direct coupling analysis (DCA), on the other hand, consider the co-evolving of all reissue pairs simultaneously and are thus able to eliminate transitivity and disentangle direct interaction from indirect interaction ^18^.

With the successful use of co-evolutionary signals within the same protein (intra-protein co-evolution) to predict protein foldings, the question arises whether inter-protein coevolution can similarly be exploited, allowing for the prediction of protein-protein interactions^13, 19–21^. For example, Cong et al. used the local statistical model mutual information (MI) (which is fast by considering each residue pair independently) and the global statistical method direct coupling analysis (DCA) (which can eliminate transitivity by considering all residue pairs comprehensively) sequentially to filter and detect protein-protein interaction (PPI) at a proteome-wide level^13^. Green et al. use a similar method as DCA called EVcomplex2 to detect PPIs in E.coli ^21^.

While direct coupling analysis has been designed to reduce transitivity (and has been used mostly within individual proteins or between proteins in direct physical contact), it does not fully solve the transitivity issue and may still occasionally report interactions between residues that are separated by relatively large physical distances. At least to some extent, this is due to the underlying biology: compensatory mutations may indeed be located at a distance, for example, due to allosteric interaction networks, ligand-mediated interactions, and so on^16, 22^. In the context of protein-protein interaction prediction, this may lead to situations where predicted protein pairs are not necessarily in direct contact - but they can be expected to at least be in the same multiprotein complex. This is often sufficient in global all-against-all protein interaction searches, which are mainly done to uncover novel functional connectivities in a given proteome, but are less concerned with the detailed physical arrangement of the newly discovered interactions.

Indirectly interacting proteins in the same complexes function concertedly to take part in various biological processes, including cell cycle regulation, differentiation, protein folding, translation, transcription, post-translational modifications, gene expression, enzyme inhibition, and antibody–antigen interactions^23^. They can be used to construct functional association networks for multiple applications such as gene prioritization, functional annotation, comparative interactomics, drug target discovery, drug repurposing, and precision medicine^24, 25^. For this purpose, the STRING database included a physical interaction score that measures the probability of two proteins being together in a trusted gold standard complex^25^. Mostly these gold standard complexes come from experimental methods. While coevolution based methods now provide a new way to detect proteins in the same complex computationally. Although we could not be sure if protein interactions detected from DCA methods are direct (direct PPI) or indirect (mediated PPI). Many computational methods could be further applied to extract direct physical interactions^19, 26^. For example, we could apply deep learning methods like AlphaFold-Multimer^27^ to investigate further if these pre-selected protein pairs directly interact. Cong et al. found in their paper that the better accuracy of direct protein interaction prediction could be achieved by down-weighting proteins that appear to coevolve with many others through a protein level average product correction (APC)^13^. One of the reasons many proteins coevolve with too many others might be due to the transitivity problem (especially when one complex exists in more than one complex). Therefore protein level APC could be a way to extract direct protein interactions further.

The effectiveness of coevolution-based methods can be restricted by three factors. Firstly, multiple sequence alignments (MSA) may be constructed with too few sequences, for example, in the case of rare gene families. Secondly, phylogenetic biases may be introduced due to an unequal taxonomic distribution of the sequences in the MSA. Thirdly, entropic biases may be an issue, in which columns of an MSA with higher entropy/variability tend to give higher MI or DCA scores than columns with low entropy^6, 28^. Various normalization methods have been proposed to alleviate this, such as average product correction (APC), which works at the residue level or even protein level and has been successfully applied in recent studies^6, 13, 17, 28, 29^.

When constructing MSAs for the detection of inter-protein coevolution, another set of challenges arises: these MSAs must be “paired,” i.e., contain two distinct proteins in each row (these are the proteins to be tested for an interaction). It must be assured that all representatives of the two proteins in the MSA share the same evolutionary trajectories to best assess the co-evolutionary signal in their sequences. This can be complicated by differential gene loss, gene duplications, and horizontal gene transfers. More specifically speaking, to build paired MSAs, researchers first have to extract orthologous proteins of each of the investigated proteins from various organisms (we refer to them as putative interologs^30^) and then concatenate them together in the same row in a paired MSA^13, 19, 31–33^. Identifying exact interologs is not trivial as most organisms contain more than one paralog for the same protein family^31, 32, 34^. Following the ortholog conjecture, paralogs are less likely to maintain the same functions than orthologs, and paralogs, after duplication, may lose or gain their interactions with other proteins^32, 35^. Several algorithms have been proposed so far to infer suitable orthologs, some of which are cumbersome and can significantly slow down the MSA construction process^36–39^. Compared with eukaryotes, prokaryotes usually show less abundant paralogy. And usually, chromosomal colocalization information in prokaryotic genomes (synteny) can help to find correct interologs, as they are frequently coded in the same operons and consequently co-transcribed^31, 36, 40, 41^. Mainly because of this reason, most coevolution-based PPI prediction studies are mainly carried out in prokaryotes, and their application to eukaryotes remains a challenge^31, 36, 40^.

Co-evolution based signals are particularly powerful when used as an ingredient in machine learning models; a prominent example of this is AlphaFold. In the latest CASP14 meeting, AlphaFold has achieved remarkable success in protein structure prediction^42, 43^. Following a similar principle, the less accurate but faster RoseTTAFold was then also proposed^44^. After the open release of AlphaFold, RoseTTAfold, and Colabfold^45^, which makes modified AlphaFold and RoseTTAFold accessible to users via Google Colab, the community found that AlphaFold type models can also be utilized to predict protein interaction after some adjustment^27, 46, 47^. The input to AlphaFold type methods to predict PPI are paired MSA data, and researchers have found optimal MSA is of pivotal importance for accurate PPI prediction, and the faster MSA generation process would significantly increase the speed of these methods^46^. Therefore, accurate and faster strategies to prepare and utilize paired MSA for the purpose of PPI are urgently needed.

To the best of our knowledge, all current coevolution-based PPI prediction approaches construct paired MSA by simply including sequences from all available bacterial genomes. However, with a quickly growing number of genomes in total, the question arises whether more is always better. First, because of the existence of paralogs which may occasionally escape proper classification by orthology assignment techniques, the inclusion of more protein sequences in paired MSAs might increase the chance of adding false interologs (paralogous protein pairs). These could confound the real co-evolutionary signals and decrease PPI detection power^33, 48^. Second, larger numbers of sequences may make it more difficult to build good MSAs given limited computational resources. Third, when paired MSAs are used to predict protein interactions, the assumption is that these interactions are conserved across all protein pairs in the MSA. However, the larger the taxonomic distances that are covered within the MSA, the more likely it is that this assumption may occasionally fail: orthologs may change interaction partners or have subtle differences in the interaction modes^16, 22^. Lastly, larger MSAs can make the preprocessing and co-evolution signal detection steps slower, which is especially problematic for proteome-wide PPI predictions.

Here, we explore strategies for making the best use of the available genome data for co-evolution protein-protein interaction prediction (Figure 1). Our focus is on interaction discovery, not on the molecular details of interactions that are already known. We find that dividing MSAs into multiple smaller MSAs (i.e., aligning separately per taxonomic clade) improves alignment quality and reduces prediction noise. By integrating the clade-wise predictions using simple machine learning, the overall predictive power is noticeably improved. Our approach can easily be executed genome-wide, and the discovered interactions can be further refined using AlphaFold.

**Fig. 1.**
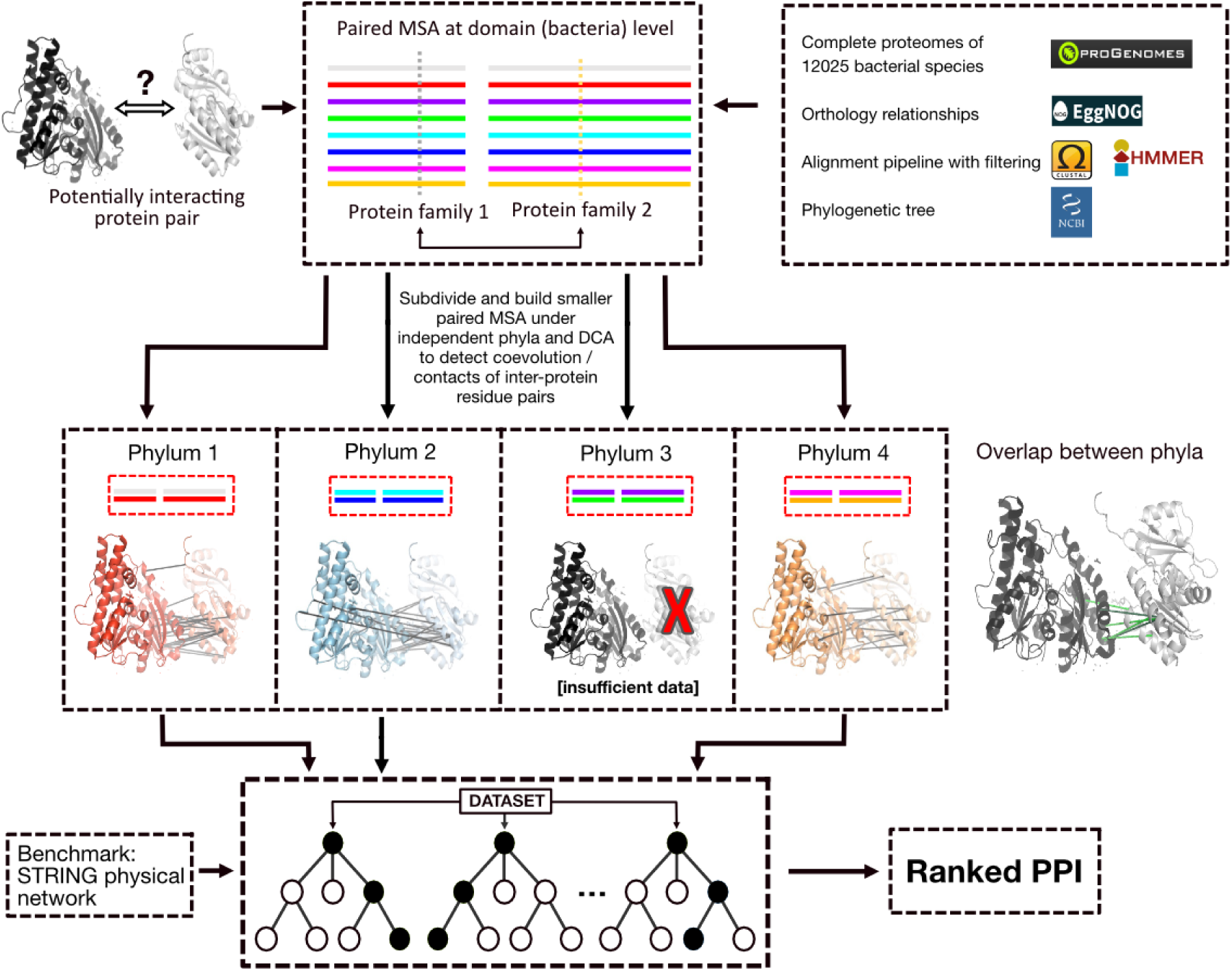
Proposed workflow of coevolution signal integration across phyla. This plot illustrates the workflow using the interacting proteins kdpC and kdpA as an example (PDB entry 5mrw^49^). Briefly, for a given protein-pair of interest, we construct paired MSAs independently in each of four phyla, and apply the DCA co-evolution algorithm to each of them separately (middle panel). Then, the top-ranking inter-protein DCA scores from each phylum are concatenated and fed to a Random Forest model (bottom panel) for an integrated evaluation and interaction prediction. The left part of the middle panel shows the positions of the top 20 inter-protein DCA scores in each phylum (all mapped to the same PDB entry). The right part of the middle panel shows the overlapped of positions detected in two phyla (light green), or three phyla (dark green).

## Results

### Alignment strategies can influence PPI prediction power

Previous work has suggested that the ability to detect co-evolutionary signals increases with the number of sequences in a given alignment. However, including too many sequences may eventually reduce alignment quality; this may also happen when including sequences that are phylogenetically too distant or whose orthology status is uncertain. Furthermore, the optimal alignment strategy may depend on the application: it may matter whether the aim is to discover novel PPIs, or to predict precise atomic arrangements within known PPIs.

To test the influence of various alignment strategies, we used known interactions in E.coli as a benchmark. First, for a given phylogenetic depth, alignments with different numbers of sequences were generated, by randomly downsampling the full alignments. This was done on paired alignments, i.e, collecting all orthologs for a given pair of proteins in E.coli, downsampling as required and then using the maximum inter-protein DCA score to predict whether the given pair might interact (or be at least be part of the same protein complex).

As expected, increasing the number of sequences in the paired alignment does lead to an increase in overall predictive power (Figure 2A). However, upon reaching about 100 sequences, the power does not improve further, but instead starts to fall again (particularly for more ‘difficult’ protein pairs at the high-recall end). Remarkably, as little as three sequences can be enough for the DCA algorithm to start picking up co-evolution and generate better-than-random PPI predictions (Figure 2A). We next tested the effects of phylogenetic (taxonomic) depth. Restricting the taxonomic level from which sequences are drawn makes the alignment process easier, since the sequences are more similar to each other, and helps to distinguish orthologs from paralogs. Indeed, as shown in Figure 2B, for a given number of sequences in the MSA, restricting their taxonomic range increases PPI prediction performance. This is concomitant with measurable improvements in alignment quality (Figure 1C): taxonomically restricted alignments contain fewer gaps and have lower column entropy. Remarkably, the optimal number of sequences remains around 100, irrespective of the taxonomic level from which the sequences are drawn.

**Fig. 2.**
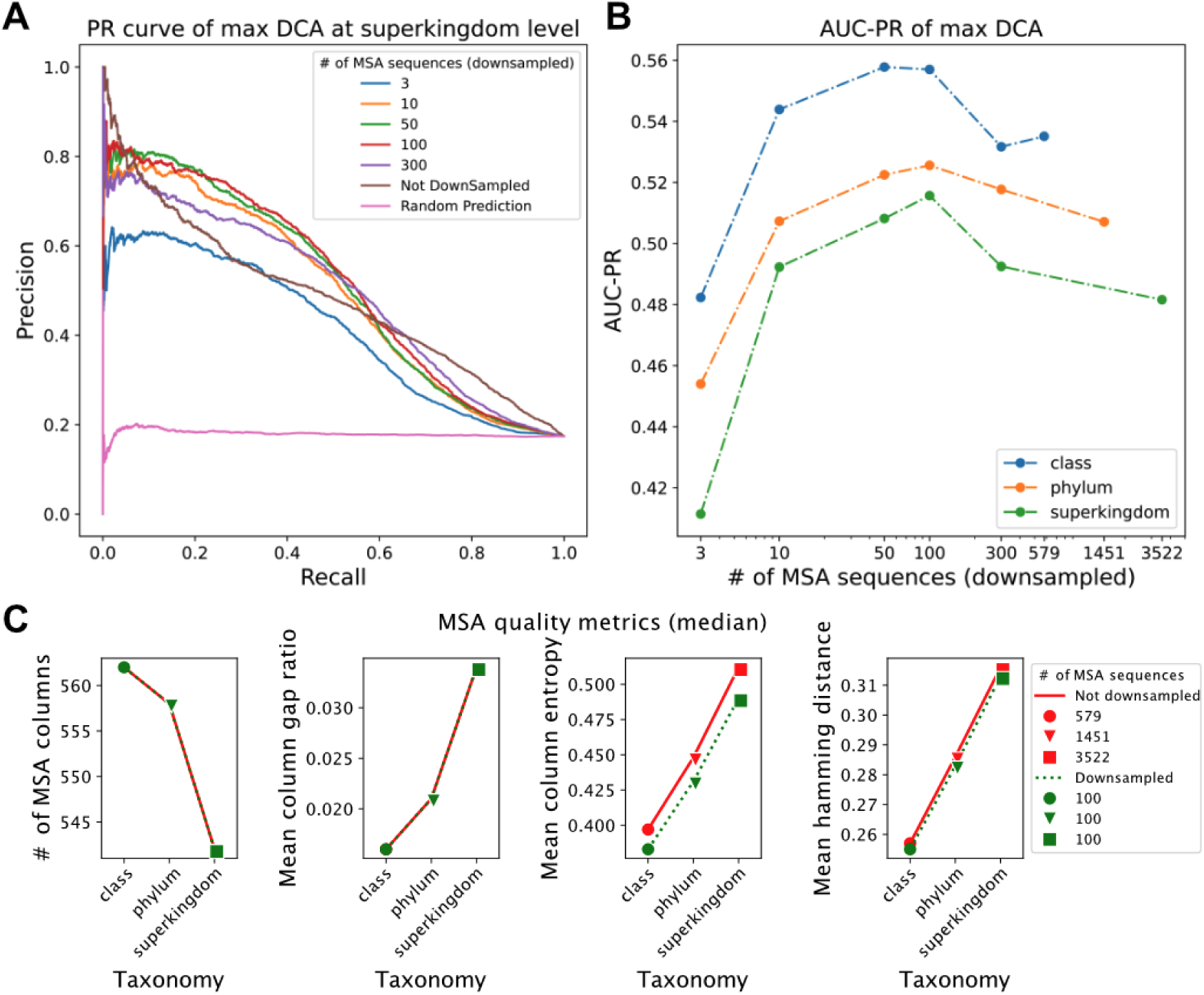
Distinct alignment strategies in coevolution-based PPI prediction. **A** PR (Precision-Recall) curve of prediction performance of varied paired MSA sizes at the superkingdom (Bacteria) level; here predictions are ranked simply according to the single highest inter-protein DCA score. The benchmark used here consists of co-complex positive PPIs and negative controls (see method). **B** AUC-PR (area under the PR curve) of prediction performance of varied paired MSA at class, phylum, and superkingdom levels. **C** Quality metrics of paired MSAs, original sizes or down-sampled (size 100).

The above observations suggest that simply including all available sequences may not be the best practice for building paired MSA for co-complex PPI prediction purposes. Given the tremendous growth in available sequences, sufficient numbers are now available to restrict the alignments taxonomically, even after preprocessing steps to remove fully redundant or otherwise problematic sequences.

### Transitivity of co-evolution signals in protein complexes

Algorithms such as DCA have previously mostly been used to study proteins in direct physical contact, but compensatory mutations may occur over fairly large distances^16, 22^. Since DCA essentially works by making observed co-evolution signals more sparse (aiming to enrich for direct couplings), its predictive power regarding longer-distance couplings in protein complexes may be different from that of shorter-distance couplings, which in turn might influence the relative merits of alignment strategies.

This is illustrated in Figure 3, using a known protein complex as an example. Three of the subunits of the high-affinity potassium pump (‘kdp’) happen to be arranged in this complex in such a way that two of them are not in direct physical contact (kdpB and kpdC); their association is mediated by the centrally-located kdpA. We built paired MSAs for all three possible protein pairs in this complex, as well as a triple MSA that included all three proteins kdpA, B, and C together. Remarkably, using the paired MSA of the two *non-contacting* proteins, the DCA algorithm nevertheless found high-scoring co-evolution signals. When including all three proteins simultaneously in the triple MSA, we found that the positions of top-ranking inter-protein residue pairs remained stable for direct protein pairs (kdpB, kdpA) and kdpA, kdpC) (Figure 2A), since in this case, both triple MSA and paired MSA contain the necessary sequences for all direct and indirect coevolution between proteins. But for the non-contacting (mediated or “bridged”) protein pair, the reported top-ranking positions of the residue pairs forming the interactions changed, indicating that there are multiple “solutions” for the DCA algorithm, which nevertheless leads to high scores. Taken together, this suggests that DCA is capable of detecting both direct and mediated protein pairs in the same complex.

**Figure 3.**
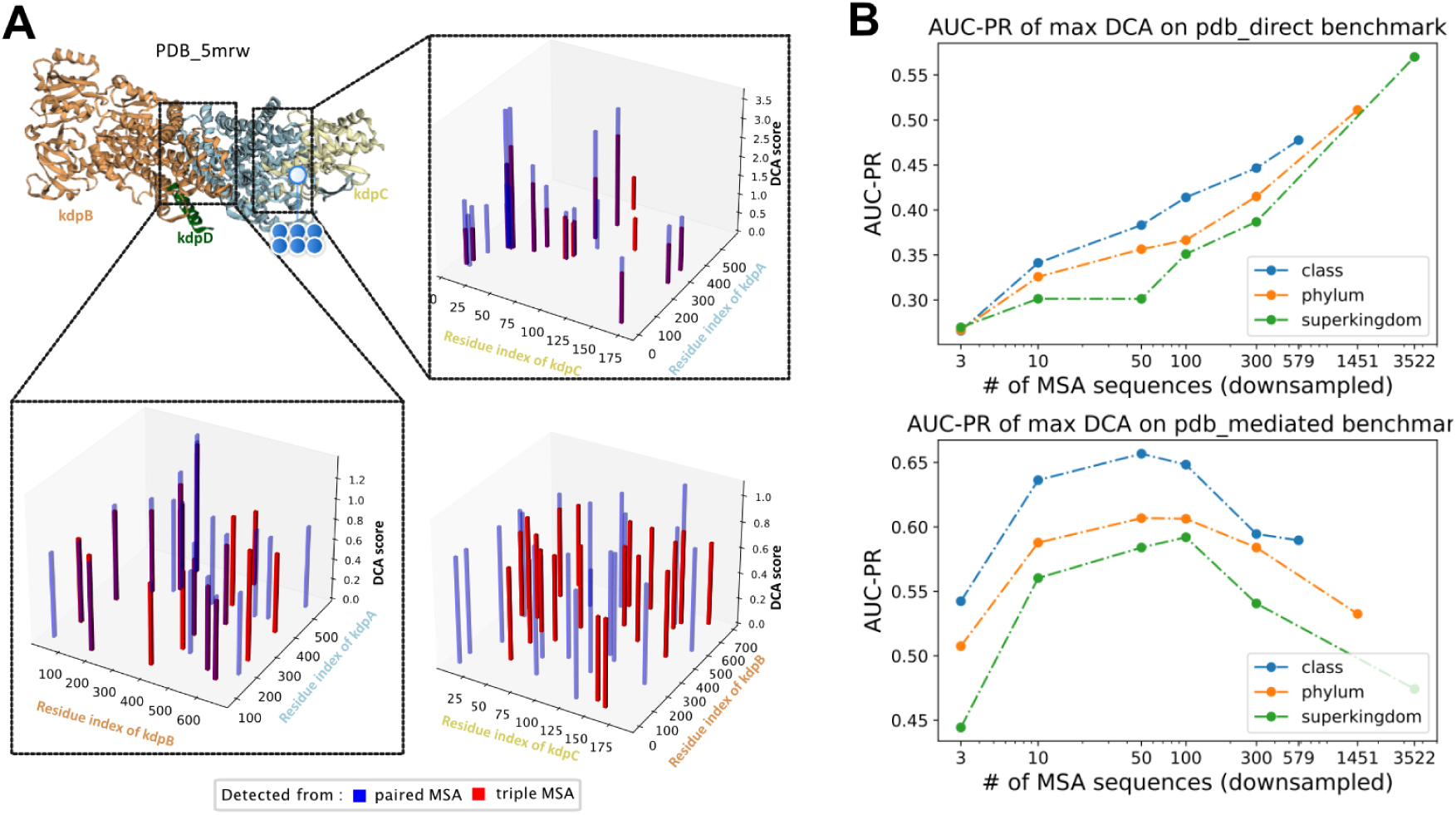
DCA performs differently on direct vs. “bridged” contacts. **A** Comparing predicted contact positions depending on whether or not other proteins are present in the MSA. At the direct interfaces (kdpA/B or kpdA/C) the presence of a third protein does not change the DCA positions or amplitudes much. In contrast, whether or not the “bridging” protein A is present between kdbB and kdpC, changes the position (but hardly the amplitudes) of DCA interactions. **B** AUC (area under PR curve) of prediction performance of one maximal inter-protein DCA score using paired MSAs at class, phylum and superkingdom levels, using direct PPI or mediated PPI benchmarks.

We systematically tested the prediction performance of DCA in both situations, using two separate benchmarks. The first is the “pdb_direct” benchmark, containing only directly binding protein pairs (see method section). The other is the “pdb_mediated” benchmark, containing protein pairs that are found in the same complex but are separated by other mediating proteins. Both benchmarks draw from the same set of non-interacting protein pairs as negatives, and both contain the same ratio of positives to negatives. We found that DCA indeed has predictive power on both benchmarks (Figure 2B). Interestingly, as we observed previously, the performances are generally better at the lower taxonomic levels, in both benchmarks. The benchmarks differ in how they rate performance of alignments with fewer sequences: For mediated contacts, relatively few sequences are sufficient, and there is an optimal number at around 100 sequences in an alignment. For direct contacts, on the other hand, more sequences generally lead to better performance, and there does not seem to be an optimum.

### Phylum-level integration of coevolution signals

While paired MSAs at narrower taxonomic clades (with fewer protein sequences) proved to be more predictive, we nevertheless wanted to find a way to use all available Bacterial protein sequences in the predictions. The intuition was that interactions should often be detectable in each Bacterial clade independently, and that integrating these predictions across the clades should increase the signal but average out the noise. We tested several ways of integrating predictions, including a simple machine learning model trained to down-weigh inconsistencies and noise.

In deciding which taxonomic rank to choose for the individual MSAs before integration, we had to strike a balance: narrow taxonomic ranges give better performance but may not always have enough genome coverage in all parts of the tree of life. As a compromise, we decided to build paired MSAs under the *<u>phylum</u>* level for integration and analysis, for the time being. In the future, considering the dramatic increase in the number of sequenced genomes, paired MSAs built under the *class* rank or ever lower taxonomic ranks could eventually lead to even better performance.

Intuitively, predictions obtained in different clades for a given protein interaction should show overlaps in terms of the positions of the interacting residue pairs. Indeed, manual spot checks showed high levels of overlap despite the paired MSAs of these orthologous protein pairs sharing no common protein sequences (see Figure 1 middle for an example). We chose four different phyla that consistently had sufficient genome coverage (*Proteobacteria*, *Firmicutes*, *Actinobacteria*, and *Bacteroidetes*). The whole pipeline is illustrated in Figure 1 and explained in detail in the Methods below. Briefly, for a given protein-pair of interest, we constructed paired MSAs independently in each of the four phyla, and applied the DCA co-evolution algorithm to each of them separately. Then, the top 5 inter-protein DCA scores from each phylum were concatenated (resulting in a vector with 20 elements) and fed to a Random Forest model tasked with classifying them into “interacting” or “non-interacting”. In case of missing data in one or more phyla, a low DCA score of -1 was entered into the vector.

For training, we again used a large benchmark containing positive and negative protein pairs in E.coli (see Methods). This benchmark was carefully balanced in terms of the evolutionary conservation of the positive and negative interaction pairs, as otherwise the machine-learning model was observed to pick up a phylogenetic cooccurrence signal from the amount and distribution of missing data from the vector (Supplementary Figure 1). Phylogenetic cooccurrence is itself a good predictor of protein-protein interactions^19^, but not of interest here. We observed that the final Random Forest model yielded a better overall performance than two simpler integration alternatives: to either rank predictions by the single, highest DCA score observed, or to apply the same Random Forest training simply on the top 20 DCA scores from a single phylum (Figure 4).

**Fig. 4.**
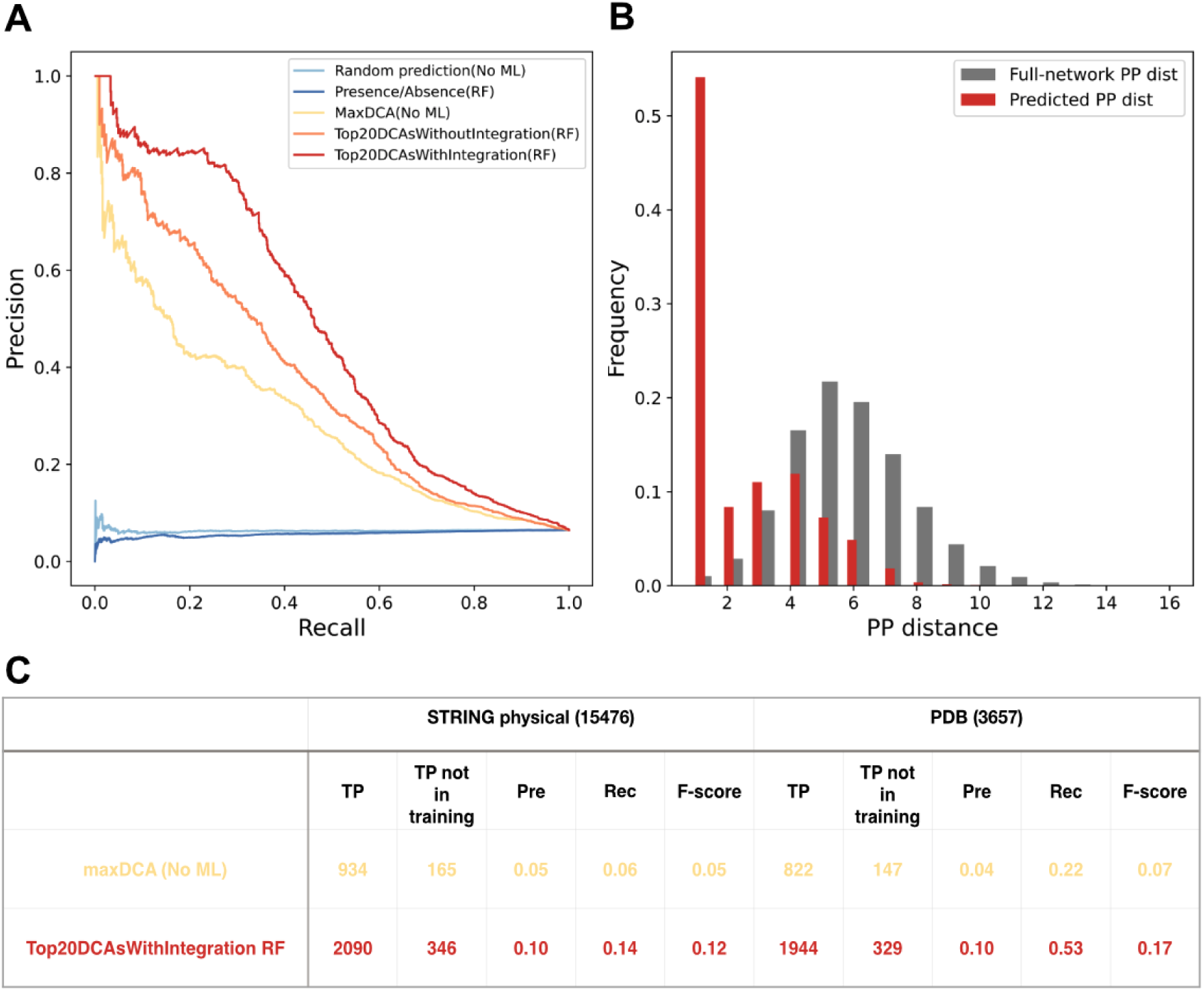
Performance of coevolution signals for co-complex PPI prediction. **A** ROC curve of prediction performance of coevolution signal integration on the refined benchmark (on the 20% testing dataset). **B.** Distance distribution (shortest paths lengths) of predicted protein pairs in the STRING physical subnetwork, comparing PPIs from the RF model to randomly paired proteins. We selected protein pairs whose predicted probabilities are more than 0.9. Among 4501 selected protein pairs, 2458 can be mapped to the STRING physical sub-network. **C.** Performance comparison of RF model trained on the integrated top 20 inter-protein DCA score from different phyla, compared to a simpler method based on only the one maximal inter-protein DCA score without any machine learning. For comparison, we selected both top 20000 PPI from two methods.

**Supplementary Fig. 1.**
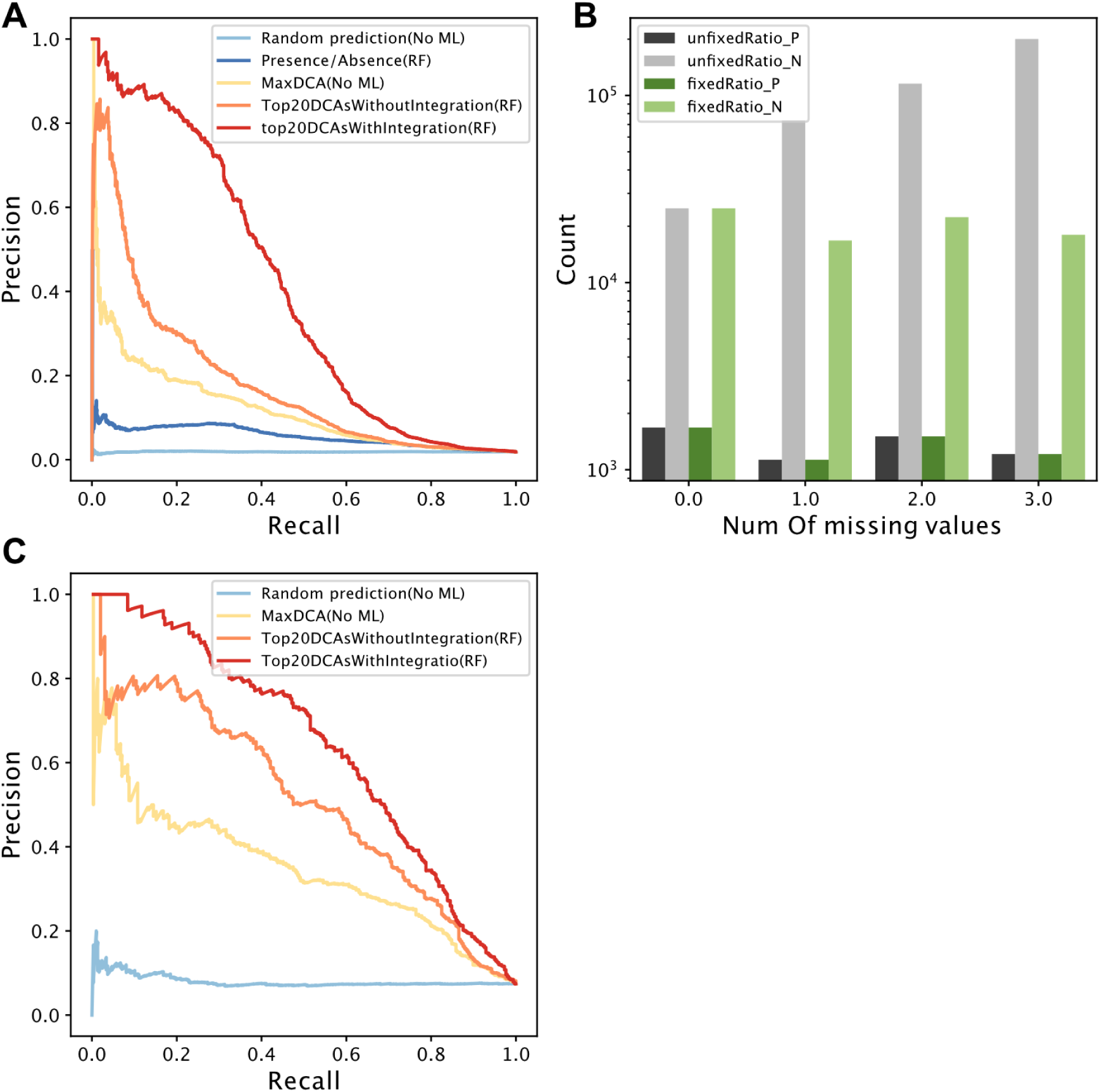
**A** ROC (receiver operating characteristic) curve of coevolution signal integration prediction performance on the initial benchmark. **B** Number of negative and positive samples on the benchmarks for coevolution signal integration before and after fixing the ratio of the number of negative and positive samples in different sample groups according to their missing values. **C** ROC curve of prediction performance of coevolution signal integration on the benchmark after removing all samples with any orthologous protein pairs in subject species.

### Proteome-wide PPI predictions

We next applied the above pipeline, including the trained Random Forest model, to all 2,269,162 protein pairs in E.coli for which sufficient, well-alignable orthologs are available. The overall performance was good, with the highest-scoring predictions achieving a precision of over 80% (at 20% recall, see Figure 4A). Also, here, the Random Forest integration over the phyla performed better than Random Forst applied to a single phylum. Furthermore, when mapping the predicted protein pairs to the physical sub-network of the STRING database, we found they tend to be in much closer proximity in the STRING network than randomly paired proteins, as expected (Figure 4B).

For these all-against-all predictions, the RF model was not re-trained, and the original training data remained as part of the benchmark. To get a lower bound on the performance for any novel, unseen interactions in E.coli, the protein pairs already included in the training dataset were removed from the prediction results as well as from the benchmark (Supplementary Figure 2). Here again, the prediction achieved > 80% precision, albeit at lower recall. This is perhaps expected: the removed interactions from the training dataset consisted of interactions that were particularly well-conserved and well-supported, and removing those left largely less-well-characterized interactions that can be assumed to be somewhat weaker and more difficult to predict.

We re-benchmarked the all-against-all results using two distinct benchmarks: the “STRING physical positive benchmark” and the “PDB positive benchmark”. These two benchmarks both reflect the co-occurrence of protein pairs in the same complex, containing both direct and mediated protein pairs, but the “STRING physical” benchmark draws from more data sources and is more comprehensive. When arbitrarily designating the top-scoring 20000 protein pairs as the “positive” predictions, it is again clear that RF integration can achieve better performance than max DCA alone (Figure 4C).

**Supplementary Figure. 2.**
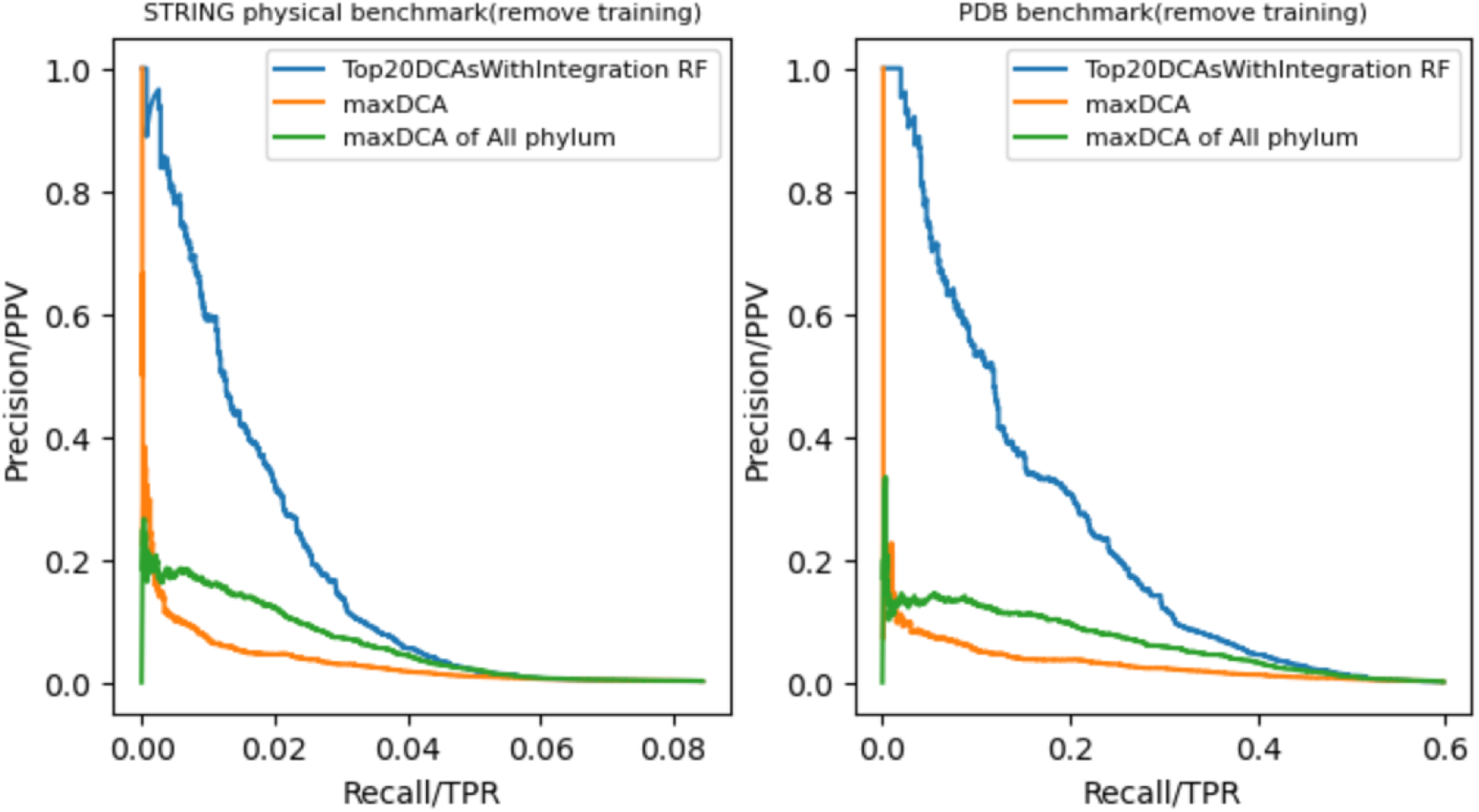
Prediction performance of coevolution signal integration at proteome level after removing training samples proportionally.

### Direct PPI detection

#### Comparison/Study with AlphaFold-Multimer

AlphaFold was originally devised to predict the fold of single proteins, but it can also be used to arrange multiple proteins in their quaternary structure in a protein complex. We tested the performance of AlphaFold-multimer ^27^ in discovering novel interactions (which is a different task than predicting the precise molecular arrangement of a known interaction). This was based on the same “pdb_direct” and “pdb_mediated” benchmarks discussed earlier (see Methods). We found that AlphaFold performs better than our DCA/clade-integration method in the pdb_direct benchmark, while our method performs better than AlphaFold on the pdb_mediated benchmark (Supplementary Figure 3). On the one hand, it is reassuring that AlphaFold can correctly discover physically interacting protein pairs. On the other hand, and somewhat counterintuitively, it tends to assign even higher contact probabilities for residues between mediated (bridged) protein pairs than direct binding protein pairs (Figure 5). AlphaFold-Multimer has been trained specifically on PDB complexes in order to predict their internal structure/contact interfaces for given PPIs. The model’s objective is not to perform global, de-novo PPI screens, and thus less attention may have been given to non-interacting, negative protein pairs in training^27, 50^.

**Fig. 5.**
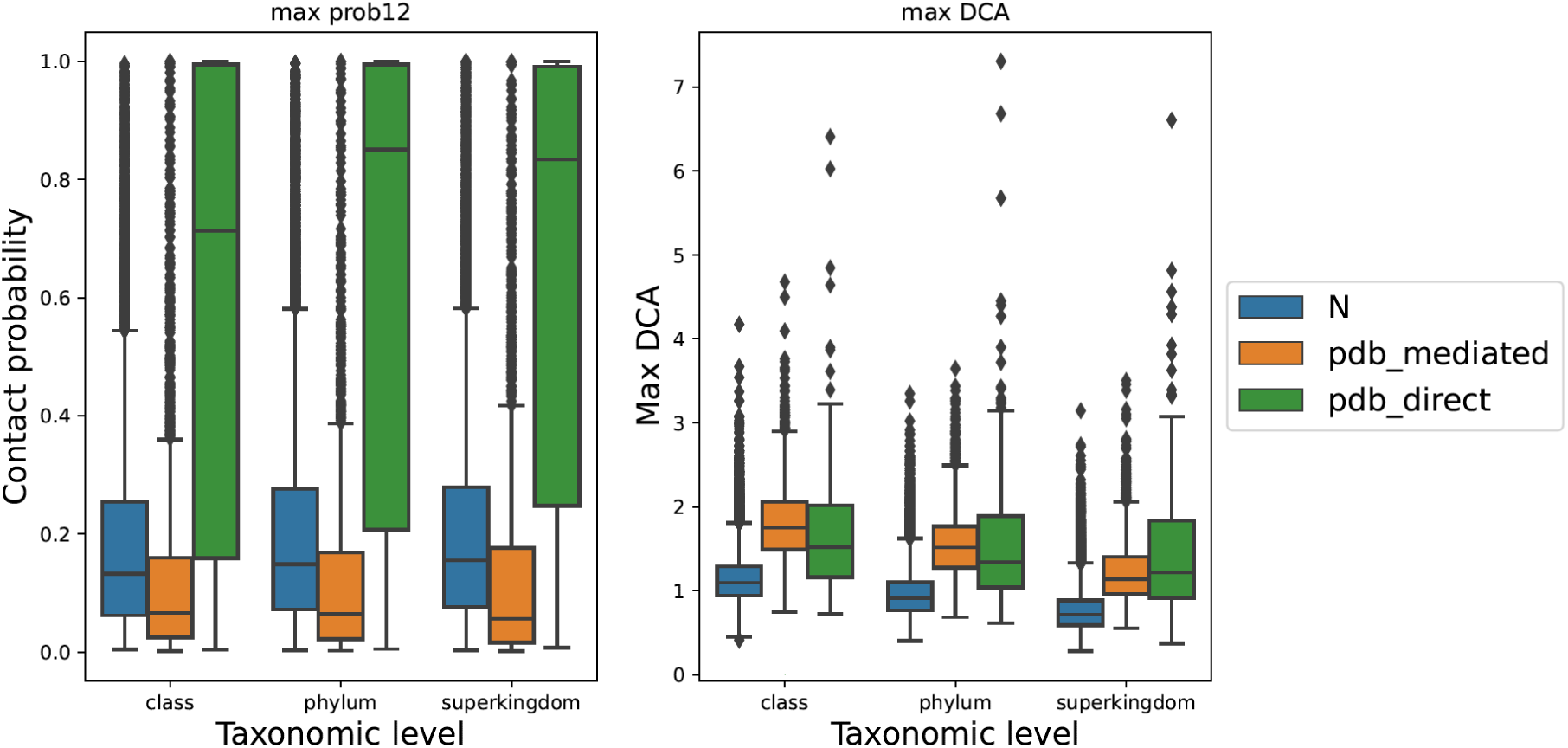
Boxplot of max in-DCA score and maximal AlphaFold contact probabilit at different protein pair groups on the common pdb_direct and pdb_mediated benchmarks at different taxonomic levels

**Supplementary Fig. 3.**
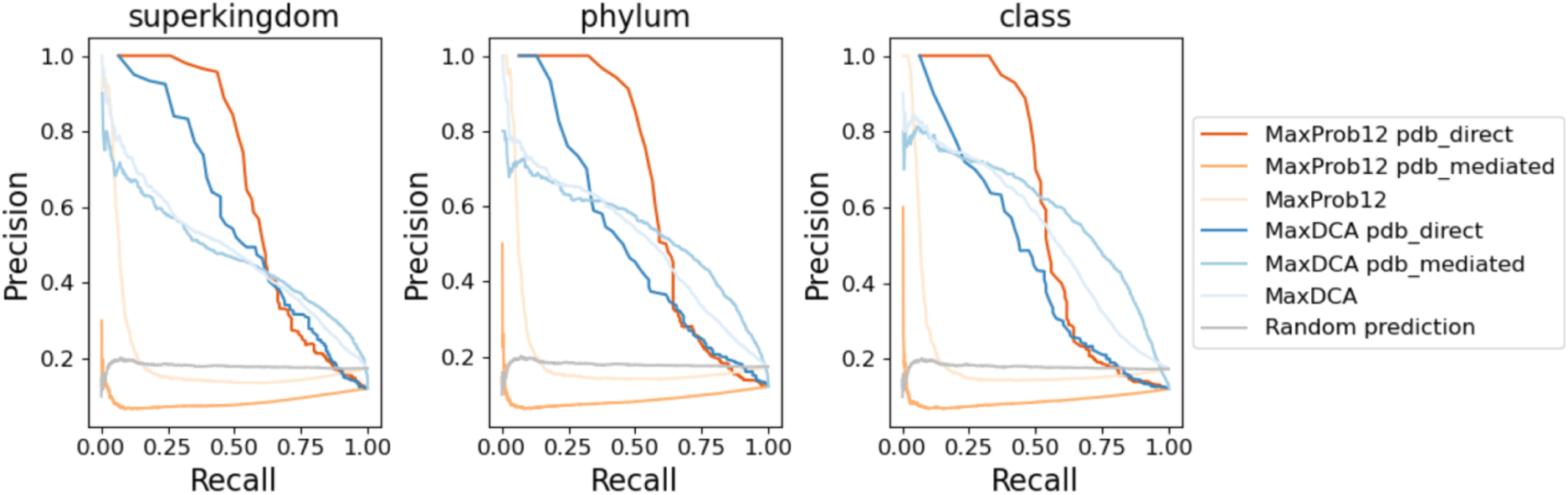
Comparison between coevolution and AlphaFold results. Precision-Recall curve of prediction performance of max coevolution signal and maximal AlphaFold contact probability on the common pdb_direct and pdb_mediated benchmarks at different taxonomical levels

#### Filtering for direct interactors

Initially, our PPI predictions from the genome-wide pipeline yielded a mix of two interaction types (direct interactions, as well as mediated/bridged interactions). From the prediction results of 2269162 protein pairs predicted by the final, best-performance RF model, we designated all 4501 PPIs that had predicted probabilities larger than 0.9 as likely enriched in direct, physical interactions. To further enrich direct PPIs, we applied the APC correction^13^ on the ranked PP list, to down-weigh proteins that appear to convolve with a large number of others. We then ran Alphafold-multimer for our top-ranking protein pairs and selected 379 with maximal inter-protein contact probabilities larger than 0.9 as our final direct PPI predictions.

We compared the performance of our pipeline with experimental high-throughput methods: one yeast two-hybrid (Y2H)^51^, and two affinity purification mass spectrometry (APMS)^52, 53^ datasets. Additionally, we compared against one computational pipeline that is also based on coevolution, from Qian’s coevolution+ ^13^. Qian et al. use MI, DCA, GREMLIN, and docking sequentially in their pipeline to discover direct PPIs. The authors also add already known protein pairs reported in experimental studies or on the same operons directly to their pipeline before the GREMLIN step.

We benchmarked results from these methods on three direct protein interaction benchmarks derived from the Y2H experiment^51^, the PDB database, and the Ecocyc database^54^, containing 277, 866, and 916 positive controls^13^. All 2,269,162 protein pairs that pass our filtering steps cover 156, 382, and 515 of them, respectively. As seen in Table 1, our combined pipeline of DCA/phylum integration plus Alphafold outperforms the experimental methods regarding precision and F1-score, with the exception of Y2H. Our pipeline has comparable or better precision than the other computational pipelines, but our recall is somewhat lower. The lower recall may be due to our more restricted genome coverage; thus, fewer protein pairs entered the final screen results (our 2,269,162 vs. their 5,433,030). Considering the future growth of sequenced genomes availability for each phylum, these restrictions may soon no longer be an issue.

**Table 1.** Comparison with other methods. Performance of experimental and coevolution screens on diverse benchmarks; this is an expanded table of Figure 1F from Qian et al.^13^. Two new benchmarks were added; the size of each benchmark is shown in parentheses. Table cells are colored by performance: dark green highlights the best-performing method, per column. F-score, harmonic mean of precision and recall; Pre, precision; Rec, recall; TP, true positives.

|  | Y2H benchmark (277) |  |  |  | Ecocyc benchmark (916) |  |  |  | Extended PDB_direct benchmark (868) |  |  |  | PDB_mediated benchmark (3243) |  |  |  | KEGG (28914) |  |  |  |
| --- | --- | --- | --- | --- | --- | --- | --- | --- | --- | --- | --- | --- | --- | --- | --- | --- | --- | --- | --- | --- |
|  | TP | Pre | Rec | F-score | TP | Pre | Rec | F-score | TP | Pre | Rec | F-score | TP | Pre | Rec | F-score | TP | Pre | Rec | F-score |
| Y2H | 69 | 3.50% | 25.10% | 6.20% | 64 | 3.30% | 7.20% | 4.50% | 52 | 2.70% | 6.80% | 3.80% | 13 | 0.70% | 0.40% | 0.50% | 84 | 4.30% | 0.30% | 0.50% |
| APMS 1 | 77 | 1.30% | 28.00% | 2.50% | 96 | 1.60% | 10.80% | 2.80% | 80 | 1.30% | 10.50% | 2.40% | 126 | 2.10% | 3.90% | 2.70% | 186 | 3.10% | 0.60% | 1.10% |
| APMS 2 | 33 | 0.30% | 12.00% | 0.50% | 177 | 1.40% | 19.80% | 2.60% | 64 | 0.50% | 8.40% | 0.90% | 10 | 0.10% | 0.30% | 0.10% | 486 | 3.80% | 1.70% | 2.30% |
| Qian Coevolution+ | 60 | 3.80% | 21.80% | 6.50% | 294 | 18.80% | 33.00% | 23.90% | 170 | 10.90% | 22.20% | 14.60% | 121 | 7.70% | 3.70% | 5.00% | 393 | 25.10% | 1.40% | 2.60% |
| Phylum integration + RF+AlphaFold | 19 | 5.00% | 6.90% | 5.80% | 69 | 18.30% | 7.70% | 10.90% | 85 | 22.50% | 11.10% | 14.90% | 14 | 3.70% | 0.40% | 0.80% | 104 | 27.50% | 0.40% | 0.70% |
| Phylum integration + RF | 3 | 0.10% | 1.10% | 0.10% | 23 | 0.50% | 2.60% | 0.90% | 71 | 1.60% | 9.30% | 2.70% | 1217 | 27.00% | 37.50% | 31.40% | 1095 | 24.30% | 3.80% | 6.60% |

Since the Y2H, APMS, and Qian’s coevolution+ pipelines are designed to detect direct PPIs only, they tend to disregard mediated PPs that are also of important value to the science community. Therefore we further checked how much better our methods can perform for mediated PPI or even on functional association benchmarks where the molecular mode of interaction is not known. For this, we constructed a PDB_mediated benchmark containing protein pairs that exist in the same PDB complex but do not bind to each other. We also constructed a functional KEGG benchmark containing protein pairs that existed in the same pathway in the KEGG database. PDB_mediated and KEGG benchmarks contain 3,243 and 28,914 true positives, and all 2,269,162 protein pairs that pass our filtering steps cover 2,954 and 16,241 of them, respectively. Our “phylum integration + RF ‘’ model has much better precision and recall than all other methods (Table 1) on the PDB_mediated benchmark, whereas, on the KEGG benchmark, it is narrowly outperformed by Alphafold and Qian et al. in terms of precision. Both methods may be somewhat overtrained, as some of the protein pairs in the benchmarks overlap with their training datasets. However, our method still performs when training data is removed (see above), albeit with less recall.

For 379 predicted direct PPs with high interaction signals according to our RF model as well as AlphaFold-Multimer, 215 are already found in the STRING database (any type of evidence considered), whereas 164 are novel protein pairs with no record in the STRING database. The full exploration of all predicted novel protein pairs lies outside the scope of this study, but a full list can be found in the supplementary file Table S1. In this paper, we only visualize three examples of these predicted protein pairs to provide a general insight (Figure 6).

**Fig. 6.**
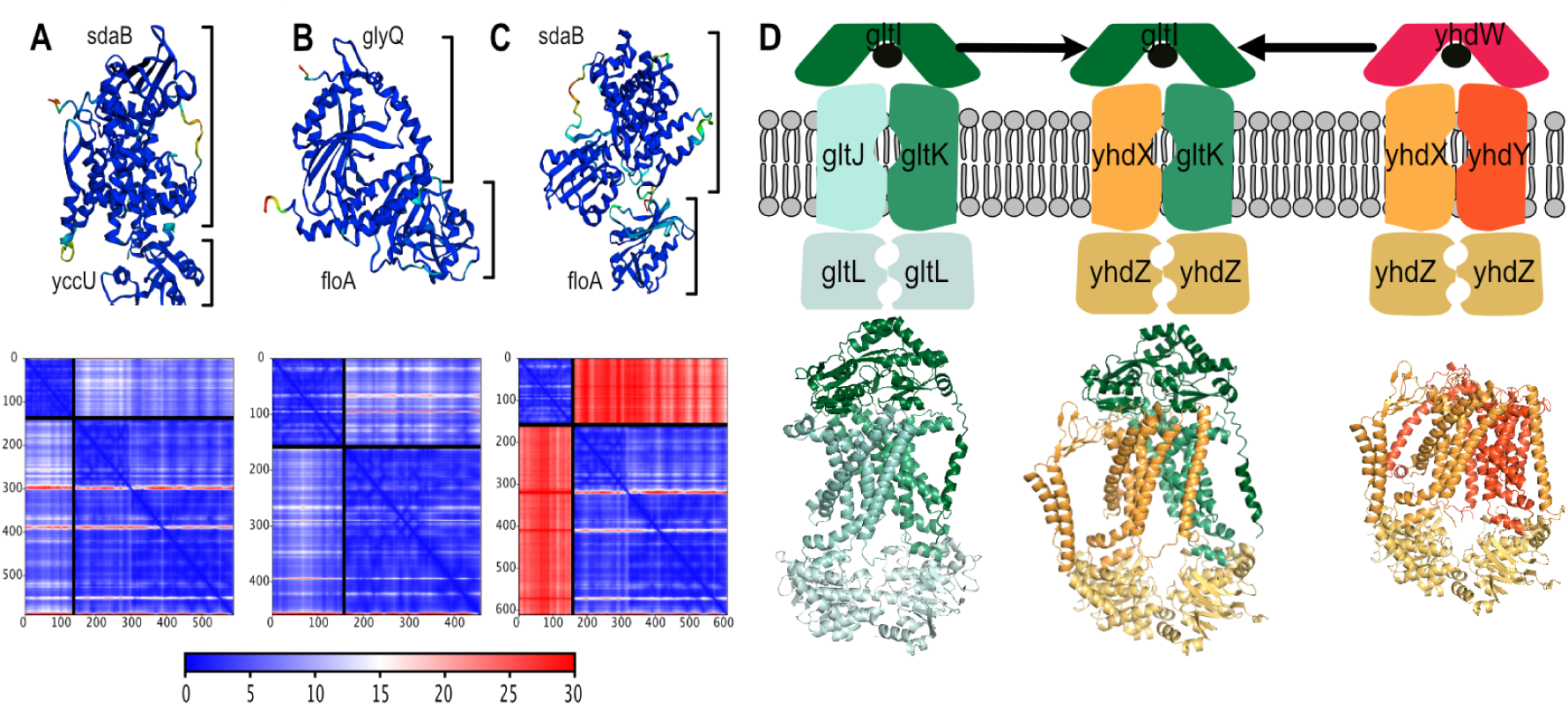
Visualization of newly predicted PPI. **A-C.** The upper panels show newly predicted pairs of interacting proteins in E.coli. The protein structures are colored by the per-residue confidence score pLDDT (blue being the maximum). The bottom panels show the predicted aligned error (PAE);there, the color at a given position (x, y) indicates AlphaFold’s expected positional error at residue x when the predicted and true structures are aligned on residue y^27^. Panel C shows a negative control; here, one protein each from panels A and B were tested for interaction. The inter-protein PAE scores are showing very high errors (red color). **D.** A novel, potential hybrid ABC transporter predicted by our pipeline. In the upper part of the figure, the left and the right illustrations show the putative glutamate/aspartate ABC transporter complex gltIJKL and putative general amino acid ABC transporter complex yhdWXYZ, respectively, from E.coli strain K-12 in the EBI Complex Portal^72^. The middle illustration shows the newly predicted hybrid ABC transporter. The bottom part of the panel shows the predicted complex structure by AlphaFold-Multimer in E.coli strain K-12 substrain MG1655. As yhdW is missing in this substrain, it is not included in the predicted structure on the right.

One protein pair that we predicted to interact without any prior known evidence is sdaB and yccU (Figure 6A). sdaB is L-serine dehydratase protein^55, 56^. yccU is so far an uncharacterized protein, but its homologs are CoA-binding proteins. It’s reported that serine can be dehydratased by sdaA/sdaB/tdcG to pyruvate, and then pyruvate can be further metabolized into Acetyl-CoA with the help of coenzyme CoA during pyruvate decarboxylation^57^. The possible involvement of both sadB and yccU (by binding to CoA) proteins in this whole metabolic pathway might support our prediction.

Another protein pair we predict to interact without any prior evidence is glyQ and folA (Figure 6B). folA catalyzes an essential reaction for a co-factor needed de novo glycine synthesis, and glyQ is the glycine tRNA synthetase. glyQ catalyzes the synthesis of glycyl-tRNA, and it needs to be covalently bound to the glycine^58, 59^. In the last step of de novo glycine synthesis, tetrahydrofolate (THF) is required, and dihydrofolate reductase folA helps to catalyze the conversion process to THF^60, 61^. The two enzymes may benefit from physical proximity, for example, via substrate channeling to facilitate rapid and efficient intermediary substrate handling^62^.

Interestedly, our model also predicted the interaction between the proteins gltK and yhdX, which are both subunits of distinct ATP-binding cassette (ABC) transporter complexes. ABC transporters complexes come in different types and functions^63^. In E.coli, they constitute the largest protein families and primarily function as importers^64, 65^. The various paralogous ABC transporters in the E.coli genome allow the cells to transport a wide range of substrates, from small molecules such as amino acids to larger compounds such as lipids and oligopeptides. ABC transporters have a characteristic architecture of two transmembrane proteins, two ATP-binding proteins, and one optional substrate-binding protein. gltIJKL is the putative glutamate/aspartate ABC transporter complex and yhdWXYZ is a putative general amino acid ABC transporter complex. The E.coli strain K-12 substrain MG1655, which is studied in this work, has both transporters, with the notable exception that the gene for yhdW is lacking from the genome.

We found several predicted interactions indicating that a “mixed” ABC transporter with subunits from both kltIJKL and yhdWXYZ might exist in E.coli. The missing yhdW could be replaced by gltI, as we also found high coevolution signal between yhdX and gltI (even though it is not in the selected top 20000 with high RF probability but ranked around the top 30000 and has AlphaFold-Multimer contact probability around 0.8). In addition, the protein yhdY could be replaced by gltK to form a hybrid ABC transporter (Figure 6D). yhdX and gltJ show some degree of sequence similarity, but their MSAs in our setup do not share common homologs. In contrast, the MSAs of yhdY and gltK do share overlapping proteins, which might inflate the co-evolution signal. We controlled for this issue by removing all shared sequences between yhdY and gltK from their paired MSA; we found the updated paired MSA still generates a high inter-protein DCA score. When dealing with large, paralogous protein families, co-evolution methods, as well as AlphaFold need to be carefully checked to avoid spurious predictions. But in this case, the scores were relatively high, and it has been reported that hybrid ABC transporters may indeed exist and remain functional^65–68^. While substrate specificities of ABC transporter are reported to be mainly determined by their substrate-binding proteins, the specificity can be further controlled by regions in the transmembrane proteins^67, 69–71^. Since both parental ABC transporters are putative amino acid transporters, our predicted hybrid form may transport amino acids as well.

## Discussion

In this study, we explored the genome-wide prediction of protein-protein interactions through co-evolution algorithms; specifically, we investigated how different alignment strategies and their concomitant differences in quality and depth of alignments could affect the predictions. We found that sufficient sequences are necessary to ensure that paired MSAs contain enough coevolution signals. But beyond a certain threshold, coevolution signals seem to become saturated. Including further protein sequences seems to introduce more noise than coevolution signals. Surprisingly, we found that for co-complex PPI prediction, better prediction performance can be achieved by building the MSAs at lower taxonomic levels with fewer protein sequences, which is in contradiction to the common practice of building paired MSAs using all protein sequences available. Further investigation suggests this contradiction may be caused by the fact that DCA performs differently for mediated vs. direct protein pairs. For direct protein pairs, the positions of direct inter-protein interactions are the same no matter how many interologs we include in the paired MSAs. In contrast, for mediated (“bridged”) protein pairs, the detected indirect coevolution positions from DCA can have some level of arbitrariness, presumably because in the absence of information about the bridging protein, DCA will have many options to choose pairs of interacting residues from a larger group of co-evolving residues^15^. For both direct as well as mediated interactions, MSAs from lower taxonomic ranks usually yield better performance when compared to MSAs from higher taxonomic ranks downsampled to the same size. This may reflect better orthology reliability, better MSA alignment quality, and fewer structural variations at lower taxonomic ranks.

Instead of building large MSAs using all bacterial sequences, we built multiple paired MSAs for a given protein pair in different taxonomic clades and then integrated coevolution signals. In this way, nearly the full set of available sequence data is used while smoothing out noise from MSAs that are too large. In this paper, we integrated coevolution signals at the phylum level; we found this to be a good compromise: it is still a fairly high taxonomic rank level, but it provides a sufficient number of sequences to allow many protein pairs to pass paired MSA alignment quality filtering steps. Due to the exponential growth of sequence data, future integration can be carried out at even lower taxonomic rank levels, such as *class*.

In our project, we used coevolution signals from all available large phyla to predict PPI, assuming that the same protein pairs have conserved interaction patterns in different clades of the tree of life. However, there might be subtle variations in how orthologous protein pairs interact in different clades. Furthermore, occasionally the process of identifying orthologs may itself be error-prone, and more so in some clades than in others^73^. In this case, using all collected signals from different clades may not be the best practice. In the future, we may need to decide further which coevolution signals to include by investigating the coevolutionary signals correlation between paired MSAs coming from different orthologous protein pairs under different phyla.

During preparation for this article, deep learning methods, especially AlphaFold from DeepMind, have succeeded greatly in protein structure prediction in CASP14^42, 43^. Following the open release of the AlphaFold paper and source code, the community also found that AlphaFold can be utilized to predict protein interactions after some adjustments^27, 47^. However, AlphaFold is still not fast enough, and perhaps also not precise enough, to be used in a genome-wide, all-against-all interaction discovery search, especially for researchers with limited computational resources. Furthermore, in its current state, it may not yet be optimal for global PPI screens due to a paucity of negative,non-interacting protein pairs in their training data^27, 50^. Our pipeline can be used as the initial screening and pre-filter step to derive promising interacting protein pairs to be fed to AlphaFold-type methods to predict the precise arrangement of the interacting partners.

## Methods

### Data source

12025 complete proteomes of all species under the *Bacteria* domain were downloaded directly from the STRING website (Version 11.5)^25^; these are in turn derived from representative bacterial proteomes in proGenomes2^74^.

Orthologous protein memberships were computed by mapping proteins from proteomes of interest to eggNOG orthologous groups (Version 5)^75^ via eggNOG-Mapper^76^.

### Paired MSA construction

The paired MSAs of protein pairs of interest from a given query species were built in two steps. In the first step, we built MSAs for single proteins. Second, we constructed paired MSAs by concatenating protein sequences from the single MSAs such that only orthologous proteins from the same species were concatenated.

For the MSA preprocessing, we adopted filter criteria similar to those used by Cong et al.^77^ to which we made some changes of our own (see below).

#### Single MSA construction

As our query organism of interest, we chose Escherichia coli str. K-12 substrain MG1655; its 4127 protein-coding genes defined the search space in which interactions were to be predicted. To remove redundant proteins, we blasted ^78^ all of these 4127 proteins with each other and treated proteins with more than 95% identity over at least 90% alignment length as redundant. For redundant proteins, we then drop shorter proteins from any further analysis.

The remaining proteins were then used as query proteins to search orthologs in other species under the desired taxonomic levels (e.g., class, phylum, and superkingdom level) via the eggNOG database of orthologous protein memberships. In eggNOG, orthology is assigned hierarchically^75^, for each taxonomic clade; when choosing orthologs for our MSA, each investigated protein’s orthologs from other species were selected at the most defined taxonomic level. If that orthologous group contains more than one protein from the subject species, then only one is selected randomly from the available orthologs. To ensure enough protein sequences in the final MSA, we kept only query proteins with orthologs in more than 1% of all the proteomes in a given clade.

Orthologous proteins of a given query protein were then used to build MSAs. A seed alignment of each query protein was first created to include its most similar orthologs via the Phmmer tool in the HMMER package^79, 80^. Three different stringency levels were used to select the seed alignment (same as Cong et. al^77^). From the most stringent criteria to the least stringent, they are (a) query coverage > 0.8 and sequence identity to the query > 0.55, (b) query coverage > 0.65 and identity > 0.4, (c) query coverage > 0.5 and identity > 0.25. For each query protein, we keep orthologs filtered by the most stringent criterion that can select over 2500 or 25% of all orthologs in the orthologous group. The actual seed alignments/MSAs of seed protein sequences were built by CLUSTAL omega software^81^. Seed alignments were then passed to hmmbuild tool from the HMMER package to build Hidden Markov Model (HMM) for query proteins. These HMMs were then used to align all the orthologous protein sequences via the hmmalign tool in the HMMER package.

Any positions that are gaps in query proteins are removed from the single MSAs since they can’t be mapped to residues in query proteins. Considering that large fractions of gaps in sequences in the given MSAs probably resulted from alignment errors or incomplete sequences, we also removed sequences containing more than 50% gaps from the single MSAs. Furthermore, MSA columns containing 50% gaps characters were removed as well, as coevolution-based algorithms may not work well for these positions. In the end, our pipeline focuses on the top-ranking inter-protein residue pairs with high coevolution signals for predicting PPIs. Therefore, even if a few positions are removed mistakenly in this step, it will not affect our final prediction too much. We also kept track of the mapping information between positions in the final MSAs and their corresponding positions in original query proteins, so that interacting residues can later be re-identified and mapped to 3D structures for analysis and visualisation.

#### MSA concatenation

After generating single MSAs for all query proteins in E. coli, we built paired MSAs for all possible query protein pairs by concatenating orthologous protein pairs from the same species/proteome at the same alignment row. Like Cong et al.^77^, we applied the HHfilter tool from HHsuite^82^ to remove highly similar sequences and sequences with too many gaps from the paired MSA (-id 90, -cov 75, -M first) to speed up the downstream computation process.

It was already reported before that the accuracy of coevolution-based methods is positively correlated with the number of sequences in MSA and negatively correlated with the square root of the MSA length. We therefore used a metric called Nf90 value (Nf90 = N90/sqrt(L)) to filter paired MSAs for downstream coevolution analysis further. Here N90 is the number of sequences in the alignment after filtering by HHfilter (-id 90, -cov 75, -M first), and L is MSA length. We used the same threshold Nf90 >=16 as Cong et al. to filter paired MSA, while our threshold should be more strict than Cong et al. as in their analysis, N90 is the number of sequences filtered by HHfilter with parameter “-id 90”. Because interacting protein pairs are more likely to co-occur in the same genome^83^, it is more likely they have high Nf90 values and thus to be included in our analysis.

### DCA computation

The aim of this article is not to compare different coevolution computation methods or to identify the most exact coevolution computation algorithm, but to find a better way to construct paired MSAs for the purpose of genome-wide PPI prediction. Therefore the relatively fast mean-field DCA algorithm, though it’s not as accurate as the pseudo-likelihood maximization algorithm, was applied on paired MSAs to compute convolution signals for the sake of computational time^16, 29, 84^. The python library used in this study was pydca: v1.0^84^.

### Interaction benchmark for single species

For the positive PPI dataset, we chose interactions from the STRING ‘Physical’ subnetwork. We filtered these on the combined physical interaction score (reflecting the likelihood of co-occurrence in the same protein complex) of more than 500^25^. Hence, positive protein pairs in this common benchmark contain both mediated and direct protein pairs.

For the negative PPI dataset, we randomly paired proteins in the positive PPI dataset. We filtered them by: 1) the interaction is not in the positive PPI dataset, but each individual protein must participate in at least one interaction in the positive dataset, 2) without any STRING score, i.e., not even a functional connection, 3) the frequency of each protein cannot be more than 60 to limit the size of the negative PPI dataset for the sake of computational time.

For both positive and negative PPI datasets, we only used protein pairs whose paired MSAs are available and valid (Nf90>=16). Despite the removal of paralogous proteins from MSAs (see section “Single MSA construction”), we occasionally observed the same protein, in a given row of the paired MSAs, to be on both sides of the pairing, i.e. to be claimed as an ortholog to both query proteins. This could indicate errors in the orthology database, or it could hint a deep and unrecognized homology between the two query proteins. Since the prediction of interactions between homologs is relatively trivial, and only few of these cases were found, we simply removed such protein pairs from our analysis entirely

Following the above-mentioned procedure, we obtained 3589, 5532, and 5910 positive protein pairs and 6039, 12960, and 19960 negative protein pairs for E.coli at class, phylum, and superkingdom levels, respectively. We did not go above the superkingdom level (Bacteria), as the inclusion of eukaryotes would increase the difficulty of finding the correct interologs required for the paired MSAs. We also did not build MSAs below the class level as there is an insufficient number of species and diversity under these levels in the STRING 11.5 database. To compare prediction performance at different taxonomic levels, we then collected and filtered the union of the above benchmarking protein pairs that could pass filtering stages at all different taxonomic levels as the final benchmark for the purpose of fair comparison. Ultimately, we obtained the same 3420 positive protein pairs and 16249 negative protein pairs for all three taxonomic levels.

Whether or not a given protein pair is predicted to interact depends on the DCA scores of its inter-protein residue pairs. To rank and benchmark the predictions, we either relied on the single, best-scoring DCA connection for a given pair, or we assessed the 20 top-scoring interactions passed through a simple machine learning model. We found that the latter approach yielded a better prediction performance; the machine learning model presumably learns that the relative strength of the 20 DCA scores compared to each other also contains information about the prediction success. We also found that increasing the number of top DCAs beyond 20 will not further improve prediction performance.

#### Down-sampling paired MSA

Down-sampled paired MSAs were produced by randomly removing alignment rows from paired MSAs built under a specific taxonomic level. No realignment was performed. If the original paired MSAs have fewer sequences than the required down-sample size, we use the original paired MSAs. This rarely happens as our maximal down-sample size is set as 300.

To assess alignment quality, we calculated the median values of the following metrics from all protein pairs in the benchmarks: number of columns in the paired MSAs (i.e., the length of the concatenated proteins after filtering); mean values of gap ratio of each column in the paired MSAs; mean values of entropy of each column in the paired MSAs; mean values of the pairwise sequence distances in the paired MSAs.

#### pdb_direct and pdb_mediated benchmarks

From the positive controls in the above benchmark, we extracted two types of interacting protein pairs. One is the *pdb_direct* group containing directly binding protein pairs (see method section “PDB benchmark”). Another one is the *pdb_mediated* group containing protein pairs in the same complex but separated by other mediating proteins. The original pdb_direct and pdb_mediated sets have 414 and 3273 protein pairs, respectively, while their overlap with this common benchmark is 154 and 2242, respectively. We then combined protein pairs from these two groups with resting negative protein pairs from the common benchmark to create two new benchmarks. For the fair comparison, we also set the same positive and negative sample ratio in the obtained pdb_direct and pdb_mediated benchmarks by randomly down-sampling the negative protein pairs. In the end, this resulted in 1116 negative samples for the pdb_direct benchmark and 16249 negative samples for the pdb_mediated benchmark.

### Coevolution signal integration across phyla

In this study, we set out to integrate coevolution signals across different phylogenetic clades to better predict whether a given protein pair of interest is interacting. Instead of a single, large MSA covering all available genomes, separate MSAs are constructed per phylum, and their coevolution signals are integrated through a machine learning model.

#### Phylum and species selection

The phyla to be chosen for separate alignments were required to have enough available genomes to build reliable MSAs. Specifically, among all 12025 representative proteomes, at present only four phyla consisting of more than 1000 species were available. They are Proteobacteria (NCBI taxonomy id: 1224), Firmicutes (NCBI taxonomy id: 1239), Actinobacteria (NCBI taxonomy id: 201174), and Bacteroidota (NCBI taxonomy id: 976) containing 4505, 2508, 1897 and 1155 genomes, respectively.

For each phylum, we chose a single representative species from which MSAs were seeded (the seeding is as described above). For the Proteobacteria phylum we chose species E.coli (NCBI taxonomy id: 511145), which we will call the query species in the context of coevolution signal integration and benchmarking. For the phyla Firmicutes, Actinobacteria, and Bacteroidota, we arbitrarily chose Paenibacillus sp. GD11(NCBI taxonomy id: 1274374), Streptacidiphilus carbonis (NCBI taxonomy id: 105422), and Bacteroides ovatus ATCC 8483 (NCBI taxonomy id: 411476), which we will call subject species in this study.

#### Orthology across phyla

For a given protein pair of interest in the query species (E.coli), orthologous protein pairs in the subject species were assigned via the eggNOG database (choosing the ‘Bacteria’ level of orthology in eggNOG). In case one of the query proteins was annotated to have multiple paralogs in a subject species, the paralog with the higher sequence similarities to the query protein was chosen (using bitscores from blastp searches, requiring an evalue cutoff of 1e-6)^78^. In the end, for each protein pair in the query species, we arrived at one best orthologous protein pair in each subject species.

#### Integrating coevolution signals from orthologous protein pairs and constructing a larger benchmark

To fully take advantage of the machine learning component, a larger benchmark was needed; it was constructed similarly to the earlier benchmark, except that we did not limit the frequency of any protein in the negative dataset. After this step, we obtained a benchmark for the query species E.coli containing 5532 positive protein pairs and 414880 negative protein pairs. This also included protein pairs in E.coli for which no orthologs were annotated in some or all of the subject species. Correspondingly, 26639, 74941, 117218 and 201614 query protein pairs have 0 or 1, 2, 3 missing orthologous protein pairs, respectively in subject species (The percentages of positive samples are 6.74%, 1.53%, 1.30%, 0.06% respectively, Supplementary Fig 1).

For a given protein pair, DCA was independently computed for all four phyla. To integrate the results, a simple strategy would be only to report the single highest-scoring inter-protein residue pair, or perhaps the average of the four highest-scoring pairs, one for each phylum. However, already within a single MSA, we observed that reporting the top-n highest scores yielded better overall performance. For the integration across the four phyla, we settled on using the top 5 best-scoring inter-protein DCA scores from each phylum (In total, each sample is a vector with 20 elements). In some cases, not all four phyla could be computed (for example, because of lacking a suitable number of orthologs). In this case, we simply concatenated the value -1 five times.

A Random Forest model was trained on this vector of top 20 DCA scores from the four phyla; it showed better performance than the a Random Forest model trained on top 20 DCA scores from a single phylum (Supplementary Figure 1A). Remarkably, we noticed that the amount of missing data (i.e. number of -1 values in the vector) appeared to be partially predictive on its own (Supplementary Figure 1B). This was confirmed by replacing the non-missing values uniformly with “+1” values - in this case, the only information available is to what extent there is missing data. Even in this case, the Machine learning model performs better than random (Supplementary Figure 1A), presumably because interacting proteins are to some extent similar in phylogenetic coverage and depth and are thus similar in the amount of missing data in their alignments.

This type of conservation information somewhat confounds potential performance improvements coming from coevolution signal integration. To prevent this type of signal from being picked up by machine learning models, we adjusted the training data such that the ratio of negative and positive pairs is the same for all levels of missing data. In the end, we got 5532 positive samples and 82131 negative samples (Supplementary Figure 1B, see Method Machine learning models). We can see from Figure 4A that the presence-absence information alone (+1 and -1) can no longer achieve a prediction performance better than random on this refined training and benchmark. Nevertheless, the conclusion that the Random Forest model trained on integrated top 20 DCA scores from four phyla can achieve better performance than the Random Forest model trained on top 20 DCA scores from a single phylum still holds.

In a separate test, in order to further demonstrate that the increased performance stems from coevolution integration and not from some other type of bias that we may have unintentionally introduced, we built another training and test set that consisted exclusively of protein pairs for which valid MSAs could be built in all four phyla. This included 1681 positive protein pairs and 24958 negative protein pairs This, as well, demonstrated performance improvements upon coevolution integration across phyla (Supplementary Figure 1C).

### Machine learning models

The main purpose of this article is to demonstrate the predictive power of coevolution integration, not to exhaustively explore which would be the most suitable machine learning setup for this. Therefore, in this study, we only assessed the simple Logistic Regression and Random Forest classifiers in Scikitlearn v.0.24.1^85^. The available data was separated into 80% training and 20% hold-out (test) data. Group K-fold (5 fold) cross-validation^86^ was applied to the training dataset. In dealing with training and testing data, we made sure that paralogous gene families were interacting only in either training and testing, but not in both. This was necessary to avoid dependent samples, and was implemented by checking for deep orthology at the eggNOG database). Grid Search was used to tune the hyper-parameters of the training models.

As a baseline control, we also predicted PPIs without machine learning models, simply by relying on inter-protein DCA scores (i.e., higher DCA scores represent high probabilities of them being positive interactions).

### Residue-level mapping of interactions

For each potentially interacting protein pair, because four separate alignments are used to assess co-evolution, a residue-level mapping between these is needed when visualizing the results. This was achieved by aligning the respective sequences from the subject species to the query species using blastp (only one sequence per alignment needed to be mapped). For visualization purposes, py3Dmol^87^ and PyMOL software was used^88^ (version 2.4.2), all results are shown in the context of the 3D structures from the query organism

### Interaction benchmarks

We constructed several distinct benchmarks for assessing different aspects of the predictions. For a general true-positive set of PPI, we chose the ‘physical’ subset of the STRING network, in E.coli K12. These were filtered by the ‘combined physical’ interaction score in STRING of more than 500 (reflecting high confidence of direct binding or at least co-occurrence in the same protein complex)^25^. It contains 15476 positive controls.

To investigate the geometries of the predicted interactions, we constructed a benchmark based on the protein data bank (PDB), containing 3657 protein pairs that are contained in the same PDB complex. This was further subdivided into two subgroups: pdb_direct (414 samples) and pdb_mediated (3243 samples). The former contains pairs for which at least 10 interacting atoms occur within 5 angstroms, while the latter contains all other pairs (usually separated by at least one bridging protein).

The broadest possible definition of an interaction is that of a ‘functional association’; for assessing this dimension, we built the KEGG functional PPI benchmark by pairing proteins that exist in the same pathways in the KEGG database (2020)^89^, resulting in 28914 protein pairs in E.coli.

### Comparisons with previous work

The ‘Y2H’ dataset is taken from a large Y2H screen in E.coli, which also contains positive PPIs compiled from manually curated databases^90, 91^ and supported by multiple publications or characterized by two independent methods^51^. The ‘extended PDB_direct’ set contains protein pairs that either interact directly with each other in the PDB^77^, or that have close homologs that interact. The ‘Ecocyc’ benchmark contains 916 protein pairs annotated to the same complexes (large complex removed); this has been extracted from the Ecocyc database^54^. The three ‘direct’ protein interaction benchmarks contain 277, 866 and 916 positive controls, respectively, and are downloaded from Cong et al.^13^.

Protein interactions according to the Y2H technology were taken from supplementary Table S2 from paper^51^. Protein interactions revealed by affinity pulldowns (APMS1 and APSM2) were taken from supplementary Table S6 of Hu et al.^53^, and from supplementary Table S2 of Babu et al.^52^. The ‘coevolution+’ list of predicted PPIs, augmented with previous knowledge), is from supplementary Table S8 of Cong et al.^77^.

### Proteome-wide PPI predictions

To identify novel PPIs at the proteome level in E.coli, we built and collected 2269162 paired MSAs at the phylum level for all possible query protein pair combinations that can pass the MSA size filtering step. All other protein pairs are treated as negative predictions. We also detected homologous protein pairs of these query protein pairs in subject species/phylum and obtained 513280, 634253, and 228993 paired MSAs in the phylum Firmicutes, Actinobacteria, and Bacteroidetes, respectively. We then computed and collected top inter-protein DCA scores from these protein pairs and applied our best-performance RF model.

When benchmarking this step, considering that some protein pairs have already been included in the training dataset of our machine learning models, we removed these pairs from our prediction results and from the benchmark. Where necessary for benchmarking, we also removed predictions to maintain the positive/negative ratio as in the specific benchmarks (Supplementary Figure 2).

### AlphaFold2-multimer computation

For our study, we downloaded the ColabFold (v1.3.0)^45^ code on 31/04/2022 and modified it in order to run it locally. Our own customized paired MSAs were then fed as inputs to ColabFold, to generate the input features that AlphaFold requires. Finally, Alphafold2-multimer (v2.2.0)^27^ is selected to predict the structure between two proteins. Alphafold2-multimer provides 5 pre-trained models. In our paper, we chose to use model 3 to save computational time. To assess the interactions between two proteins, we extracted both inter-protein residue distances (between C alphas) and contact probability of Cbeta-Cbeta under 12 Å as mentioned in paper^47^.

### The final set of predictions

From the list of 2269162 protein pairs predicted by the final best performance RF model, we selected 4501 PPIs with predicted probabilities larger than 0.9 as our final physical PPI predictions.

To further characterize novel, directly contacting protein pairs, we applied Alaphfold-Multimer to filter further results obtained from our best-performance RF model. Additionally, as Cong et al. found in their study that a better accuracy of direct protein interaction prediction could be achieved by down-weighing proteins that appear to coevolve with many others through the protein level average product correction (APC)^13^, we also ranked the PP list selected by predicted probabilities from our best performance RF model after performing protein level APC using all 2269162 protein pairs. Then, from the ranked PP list, we ran Alphafold-multimer for the top 20000 protein pairs and selected 379 with maximal inter-protein contact probabilities larger than 0.9 as the final direct PPI predictions (Supplementary Table S1).

## Acknowledgments

We would like to thank current and former members of the von Mering group, especially Dean Sumner, for valuable discussions and input.

## Authors’ contributions

TF, DS, and CVM designed the research, analyzed the data, and wrote the article. TF implemented the method. RH helped to compare various DCA algorithms. All authors read and approved the final manuscript.

## Ethics approval and consent to participate

Not applicable

## Consent for publication

Not applicable

## Availability of data and materials

Proteome sequence data and orthology information can be downloaded from STRING and eggNOG websites respectively. The code to reproduce the results in this study is available at https://github.com/TaoDFang/PPI_Prediction_byCoevolution.

## Supplementary Information

Table S1 Predicted direct PPs with high interaction signals according to our RF model as well as AlphaFold-Multimer,

## References

1. Lesk, A. M. & Chothia, C. How different amino acid sequences determine similar protein structures: The structure and evolutionary dynamics of the globins. J. Mol. Biol. 136, 225–270 (1980).

2. Marsh, J. A. & Teichmann, S. A. Parallel dynamics and evolution: Protein conformational fluctuations and assembly reflect evolutionary changes in sequence and structure: Prospects & Overviews. BioEssays 36, 209–218 (2014).

3. Haney, P. J. et al. Thermal adaptation analyzed by comparison of protein sequences from mesophilic and extremely thermophilic *Methanococcus* species. Proc. Natl. Acad. Sci. 96, 3578–3583 (1999).

4. Pál, C., Papp, B. & Lercher, M. J. An integrated view of protein evolution. Nat. Rev. Genet. 7, 337–348 (2006).

5. Brininger, C., Spradlin, S., Cobani, L. & Evilia, C. The more adaptive to change, the more likely you are to survive: Protein adaptation in extremophiles. Semin. Cell Dev. Biol. 84, 158–169 (2018).

6. Dunn, S. D., Wahl, L. M. & Gloor, G. B. Mutual information without the influence of phylogeny or entropy dramatically improves residue contact prediction. Bioinformatics 24, 333–340 (2008).

7. Buslje, C. M., Santos, J., Delfino, J. M. & Nielsen, M. Correction for phylogeny, small number of observations and data redundancy improves the identification of coevolving amino acid pairs using mutual information. Bioinformatics 25, 1125–1131 (2009).

8. Koehl, P. & Levitt, M. Sequence Variations within Protein Families are Linearly Related to Structural Variations. J. Mol. Biol. 323, 551–562 (2002).

9. Gloor, G. B., Martin, L. C., Wahl, L. M. & Dunn, S. D. Mutual Information in Protein Multiple Sequence Alignments Reveals Two Classes of Coevolving Positions ^†^. Biochemistry 44, 7156–7165 (2005).

10. Shackelford, G. & Karplus, K. Contact prediction using mutual information and neural nets. Proteins Struct. Funct. Bioinforma. 69, 159–164 (2007).

11. Jones, D. T., Buchan, D. W. A., Cozzetto, D. & Pontil, M. PSICOV: precise structural contact prediction using sparse inverse covariance estimation on large multiple sequence alignments. Bioinformatics 28, 184–190 (2012).

12. Cocco, S., Feinauer, C., Figliuzzi, M., Monasson, R. & Weigt, M. Inverse statistical physics of protein sequences: a key issues review. Rep. Prog. Phys. 81, 032601 (2018).

13. Cong, Q., Anishchenko, I., Ovchinnikov, S. & Baker, D. Protein interaction networks revealed by proteome coevolution. Science 365, 185 (2019).

14. Marks, D. S., Hopf, T. A. & Sander, C. Protein structure prediction from sequence variation. Nat. Biotechnol. 30, 1072–1080 (2012).

15. Weigt, M., White, R. A., Szurmant, H., Hoch, J. A. & Hwa, T. Identification of direct residue contacts in protein-protein interaction by message passing. Proc. Natl. Acad. Sci. 106, 67–72 (2009).

16. Morcos, F., et al. Direct-coupling analysis of residue coevolution captures native contacts across many protein families. Proc. Natl. Acad. Sci. 108, E1293–E1301 (2011).

17. Kamisetty, H., Ovchinnikov, S. & Baker, D. Assessing the utility of coevolution-based residue-residue contact predictions in a sequence- and structure-rich era. Proc. Natl. Acad. Sci. 110, 15674–15679 (2013).

18. Tetchner, S., Kosciolek, T. & Jones, D. T. Opportunities and limitations in applying coevolution-derived contacts to protein structure prediction. Bio-Algorithms Med-Syst. 10, 243–254 (2014).

19. Keskin, O., Tuncbag, N. & Gursoy, A. Predicting Protein–Protein Interactions from the Molecular to the Proteome Level. Chem. Rev. 116, 4884–4909 (2016).

20. de Juan, D., Pazos, F. & Valencia, A. Emerging methods in protein co-evolution. Nat. Rev. Genet. 14, 249–261 (2013).

21. Green, A. G. et al. Large-scale discovery of protein interactions at residue resolution using co-evolution calculated from genomic sequences. Nat. Commun. 12, 1396 (2021).

22. Anishchenko, I., Ovchinnikov, S., Kamisetty, H. & Baker, D. Origins of coevolution between residues distant in protein 3D structures. Proc. Natl. Acad. Sci. 114, 9122–9127 (2017).

23. Zahiri, J. et al. Protein complex prediction: A survey. Genomics 112, 174–183 (2020).

24. Guala, D., Ogris, C., Müller, N. & Sonnhammer, E. L. L. Genome-wide functional association networks: background, data & state-of-the-art resources. Brief. Bioinform. 21, 1224–1237 (2020).

25. Szklarczyk, D. et al. The STRING database in 2021: customizable protein–protein networks, and functional characterization of user-uploaded gene/measurement sets. Nucleic Acids Res. 49, D605–D612 (2021).

26. Laine, E., Eismann, S., Elofsson, A. & Grudinin, S. Protein sequence-to-structure learning: Is this the end(-to-end revolution)? Proteins Struct. Funct. Bioinforma. 89, 1770–1786 (2021).

27. Evans, R., et al. Protein complex prediction with AlphaFold-Multimer. http://biorxiv.org/lookup/doi/10.1101/2021.10.04.463034 (2021) doi:10.1101/2021.10.04.463034.

28. Vorberg, S., Seemayer, S. & Söding, J. Synthetic protein alignments by CCMgen quantify noise in residue-residue contact prediction. PLOS Comput. Biol. 14, e1006526 (2018).

29. Ekeberg, M., Lövkvist, C., Lan, Y., Weigt, M. & Aurell, E. Improved contact prediction in proteins: Using pseudolikelihoods to infer Potts models. *Phys*. Rev. E 87, 012707 (2013).

30. Yu, H. et al. Annotation Transfer Between Genomes: Protein–Protein Interologs and Protein–DNA Regulogs. Genome Res. 14, 1107–1118 (2004).

31. Ovchinnikov, S., Kamisetty, H. & Baker, D. Robust and accurate prediction of residue–residue interactions across protein interfaces using evolutionary information. eLife 3, e02030 (2014).

32. Feinauer, C., Szurmant, H., Weigt, M. & Pagnani, A. Inter-Protein Sequence Co-Evolution Predicts Known Physical Interactions in Bacterial Ribosomes and the Trp Operon. PLOS ONE 11, e0149166 (2016).

33. Szurmant, H. & Weigt, M. Inter-residue, inter-protein and inter-family coevolution: bridging the scales. Curr. Opin. Struct. Biol. 50, 26–32 (2018).

34. Wu, F. Y. The Potts model. Rev. Mod. Phys. 54, 235–268 (1982).

35. Altenhoff, A. M., Studer, R. A., Robinson-Rechavi, M. & Dessimoz, C. Resolving the Ortholog Conjecture: Orthologs Tend to Be Weakly, but Significantly, More Similar in Function than Paralogs. PLoS Comput. Biol. 8, e1002514 (2012).

36. Gueudré, T., Baldassi, C., Zamparo, M., Weigt, M. & Pagnani, A. Simultaneous identification of specifically interacting paralogs and interprotein contacts by direct coupling analysis. Proc. Natl. Acad. Sci. 113, 12186–12191 (2016).

37. Bitbol, A.-F. Inferring interaction partners from protein sequences using mutual information. PLOS Comput. Biol. 14, e1006401 (2018).

38. Marmier, G., Weigt, M. & Bitbol, A.-F. Phylogenetic correlations can suffice to infer protein partners from sequences. PLOS Comput. Biol. 15, e1007179 (2019).

39. Bitbol, A.-F., Dwyer, R. S., Colwell, L. J. & Wingreen, N. S. Inferring interaction partners from protein sequences. Proc. Natl. Acad. Sci. 113, 12180–12185 (2016).

40. Rodriguez-Rivas, J., Marsili, S., Juan, D. & Valencia, A. Conservation of coevolving protein interfaces bridges prokaryote–eukaryote homologies in the twilight zone. Proc. Natl. Acad. Sci. 113, 15018–15023 (2016).

41. Ren, Q. & Paulsen, I. T. Comparative Analyses of Fundamental Differences in Membrane Transport Capabilities in Prokaryotes and Eukaryotes. PLoS Comput. Biol. 1, e27 (2005).

42. Jumper, J. et al. Highly accurate protein structure prediction with AlphaFold. Nature 596, 583–589 (2021).

43. Tunyasuvunakool, K. et al. Highly accurate protein structure prediction for the human proteome. Nature 596, 590–596 (2021).

44. Baek, M. et al. Accurate prediction of protein structures and interactions using a three-track neural network. Science 373, 871–876 (2021).

45. Mirdita, M. et al. ColabFold: making protein folding accessible to all. Nat. Methods (2022) doi:10.1038/s41592-022-01488-1.

46. Bryant, P., Pozzati, G. & Elofsson, A. Improved prediction of protein-protein interactions using AlphaFold2. Nat. Commun. 13, 1265 (2022).

47. Humphreys, I. R. et al. Computed structures of core eukaryotic protein complexes. Science 374, eabm4805 (2021).

48. Bryant, P., Pozzati, G. & Elofsson, A. Improved prediction of protein-protein interactions using AlphaFold2. http://biorxiv.org/lookup/doi/10.1101/2021.09.15.460468 (2021) doi:10.1101/2021.09.15.460468.

49. Huang, C.-S., Pedersen, B. P. & Stokes, D. L. Crystal structure of the potassium-importing KdpFABC membrane complex. Nature 546, 681–685 (2017).

50. Gao, M., Nakajima An, D., Parks, J. M. & Skolnick, J. AF2Complex predicts direct physical interactions in multimeric proteins with deep learning. Nat. Commun. 13, 1744 (2022).

51. Rajagopala, S. V. et al. The binary protein-protein interaction landscape of Escherichia coli. Nat. Biotechnol. 32, 285–290 (2014).

52. Babu, M. et al. Global landscape of cell envelope protein complexes in Escherichia coli. Nat. Biotechnol. 36, 103–112 (2018).

53. Hu, P. et al. Global Functional Atlas of Escherichia coli Encompassing Previously Uncharacterized Proteins. PLoS Biol. 7, e1000096 (2009).

54. Keseler, I. M. et al. EcoCyc: a comprehensive database of Escherichia coli biology. Nucleic Acids Res. 39, D583–D590 (2011).

55. Mann, S. et al. Isolation, Characterization and Biosafety Evaluation of Lactobacillus Fermentum OK with Potential Oral Probiotic Properties. Probiotics Antimicrob. Proteins 13, 1363–1386 (2021).

56. Shao, Z. & Newman, E. B. Sequencing and characterization of the sdaB gene from Escherichia coli K-12. Eur. J. Biochem. 212, 777–784 (1993).

57. Burman, J. D., Stevenson, C. E., Sawers, R. G. & Lawson, D. M. The crystal structure of Escherichia coli TdcF, a member of the highly conserved YjgF/YER057c/UK114 family. BMC Struct. Biol. 7, 30 (2007).

58. Freist, W., Logan, D. T. & Gauss, D. H. Glycyl-tRNA synthetase. Biol. Chem. Hoppe. Seyler 377, 343–356 (1996).

59. Chen, X. et al. DCEO Biotechnology: Tools To Design, Construct, Evaluate, and Optimize the Metabolic Pathway for Biosynthesis of Chemicals. Chem. Rev. 118, 4–72 (2018).

60. Salusjärvi, L., Havukainen, S., Koivistoinen, O. & Toivari, M. Biotechnological production of glycolic acid and ethylene glycol: current state and perspectives. Appl. Microbiol. Biotechnol. 103, 2525–2535 (2019).

61. Härtel, T. et al. Characterization of Central Carbon Metabolism of Streptococcus pneumoniae by Isotopologue Profiling. J. Biol. Chem. 287, 4260–4274 (2012).

62. Huang, X., Holden, H. M. & Raushel, F. M. Channeling of Substrates and Intermediates in Enzyme-Catalyzed Reactions. Annu. Rev. Biochem. 70, 149–180 (2001).

63. Srikant, S. Evolutionary history of ATP-binding cassette proteins. FEBS Lett. 594, 3882–3897 (2020).

64. Rees, D. C., Johnson, E. & Lewinson, O. ABC transporters: the power to change. Nat. Rev. Mol. Cell Biol. 10, 218–227 (2009).

65. Moussatova, A., Kandt, C., O’Mara, M. L. & Tieleman, D. P. ATP-binding cassette transporters in Escherichia coli. Biochim. Biophys. Acta BBA - Biomembr. 1778, 1757–1771 (2008).

66. Silver, R. P., Prior, K., Nsahlai, C. & Wright, L. F. ABC transporters and the export of capsular polysaccharides from Gram-negative bacteria. Res. Microbiol. 152, 357–364 (2001).

67. Teichmann, L. et al. From substrate specificity to promiscuity: hybrid ABC transporters for osmoprotectants: Hybrid osmolyte ABC transporters. Mol. Microbiol. 104, 761–780 (2017).

68. Yang, D. C. et al. An ATP-binding cassette transporter-like complex governs cell-wall hydrolysis at the bacterial cytokinetic ring. Proc. Natl. Acad. Sci. 108, (2011).

69. Yu, J., Ge, J., Heuveling, J., Schneider, E. & Yang, M. Structural basis for substrate specificity of an amino acid ABC transporter. Proc. Natl. Acad. Sci. 112, 5243–5248 (2015).

70. Oldham, M. L., Khare, D., Quiocho, F. A., Davidson, A. L. & Chen, J. Crystal structure of a catalytic intermediate of the maltose transporter. Nature 450, 515–521 (2007).

71. Heuveling, J., Landmesser, H. & Schneider, E. One Intact Transmembrane Substrate Binding Site Is Sufficient for the Function of the Homodimeric Type I ATP-Binding Cassette Importer for Positively Charged Amino Acids Art(MP) _2_ of Geobacillus stearothermophilus. J. Bacteriol. 200, (2018).

72. Meldal, B. H. M. et al. Complex Portal 2022: new curation frontiers. Nucleic Acids Res. 50, D578–D586 (2022).

73. Lewis, A. C. F., Jones, N. S., Porter, M. A. & Deane, C. M. What Evidence Is There for the Homology of Protein-Protein Interactions? PLoS Comput. Biol. 8, e1002645 (2012).

74. Mende, D. R. et al. proGenomes2: an improved database for accurate and consistent habitat, taxonomic and functional annotations of prokaryotic genomes. Nucleic Acids Res. gkz1002 (2019) doi:10.1093/nar/gkz1002.

75. Huerta-Cepas, J. et al. eggNOG 5.0: a hierarchical, functionally and phylogenetically annotated orthology resource based on 5090 organisms and 2502 viruses. Nucleic Acids Res. 47, D309–D314 (2019).

76. Huerta-Cepas, J. et al. Fast Genome-Wide Functional Annotation through Orthology Assignment by eggNOG-Mapper. Mol. Biol. Evol. 34, 2115–2122 (2017).

77. Cong, Q., Anishchenko, I., Ovchinnikov, S. & Baker, D. Protein interaction networks revealed by proteome coevolution. Science 365, 185–189 (2019).

78. Camacho, C. et al. BLAST+: architecture and applications. BMC Bioinformatics 10, 421 (2009).

79. Eddy, S. R. Multiple alignment using hidden Markov models. Proc. Int. Conf. Intell. Syst. Mol. Biol. 3, 114–120 (1995).

80. Eddy, S. R. Accelerated Profile HMM Searches. PLoS Comput. Biol. 7, e1002195 (2011).

81. Sievers, F. et al. Fast, scalable generation of high-quality protein multiple sequence alignments using Clustal Omega. Mol. Syst. Biol. 7, 539 (2011).

82. Remmert, M., Biegert, A., Hauser, A. & Söding, J. HHblits: lightning-fast iterative protein sequence searching by HMM-HMM alignment. Nat. Methods 9, 173–175 (2012).

83. Pellegrini, M. Using Phylogenetic Profiles to Predict Functional Relationships. in Bacterial Molecular Networks (eds. van Helden, J., Toussaint, A. & Thieffry, D.) vol. 804 167–177 (Springer New York, 2012).

84. Zerihun, M. B., Pucci, F., Peter, E. K. & Schug, A. pydca v1.0: a comprehensive software for Direct Coupling Analysis of RNA and Protein Sequences. http://biorxiv.org/lookup/doi/10.1101/805523 (2019) doi:10.1101/805523.

85. Pedregosa, F. et al. Scikit-learn: Machine Learning in Python. ArXiv12010490 Cs (2018).

86. Whalen, S., Schreiber, J., Noble, W. S. & Pollard, K. S. Navigating the pitfalls of applying machine learning in genomics. Nat. Rev. Genet. (2021) doi:10.1038/s41576-021-00434-9.

87. Rego, N. & Koes, D. 3Dmol.js: molecular visualization with WebGL. Bioinformatics 31, 1322–1324 (2015).

88. PyMOL | pymol.org. https://pymol.org/2/.

89. Kanehisa, M., Furumichi, M., Sato, Y., Kawashima, M. & Ishiguro-Watanabe, M. KEGG for taxonomy-based analysis of pathways and genomes. Nucleic Acids Res. 51, D587–D592 (2023).

90. Goll, J. et al. MPIDB: the microbial protein interaction database. Bioinformatics 24, 1743–1744 (2008).

91. Rajagopala, S. V. et al. MPI-LIT: a literature-curated dataset of microbial binary protein--protein interactions. Bioinformatics 24, 2622–2627 (2008).

